# A Systematic Evaluation of the Computational Tools for Ligand-receptor-based Cell-Cell Interaction Inference

**DOI:** 10.1101/2022.04.05.487237

**Authors:** Saidi Wang, Hansi Zheng, James S. Choi, Jae K. Lee, Xiaoman Li, Haiyan Hu

## Abstract

Cell-cell interactions (CCIs) are essential for multicellular organisms to coordinate biological processes and functions. Many molecules and signaling processes can mediate CCIs. One classical type of CCI mediator is the interaction between secreted ligands and cell surface receptors, i.e., ligand-receptor (LR) interaction. With the recent development of single-cell technologies, a large amount of single-cell RNA Sequencing (scRNA-Seq) data has become widely available. This data availability motivated the single-cell-resolution study of CCIs, particularly LR-based CCIs. Dozens of computational methods and tools have been developed to predict CCIs by identifying LR-based CCIs. Many of these tools have been theoretically reviewed. However, there is little study on current LR-based CCI prediction tools regarding their performance and running results on public scRNA-Seq datasets. In this work, to fill this gap, we tested and compared nine of the most recent computational tools for LR-based CCI prediction. We used fifteen mouse scRNA-Seq samples that correspond to nearly 100K single cells under different experimental conditions for testing and comparison. Besides briefing the methodology used in these nine tools, we summarized the similarities and differences of these tools in terms of both LR prediction and CCI inference between cell types. We provided insight into using these tools to make meaningful discoveries in understanding cell communications.

## Introduction

Cell-Cell interactions (CCIs) are essential for multicellular organisms to develop tissue structure and regulate individual cell processes [1-4]. They also contribute to maintaining intercellular relationships and coordinating diverse biological processes, such as development, differentiation and inflammation [5-7]. A CCI occurs when one cell called the sender cell transmits information via signaling molecules to another cell called the receiver cell [8]. Various signaling molecules such as ions, metabolites, integrins, receptors, junction proteins, structural proteins, ligands and secreted proteins of the extracellular matrix are involved in CCIs [9, 10]. A typical signaling cascade begins with a single key event, such as ligand-receptor (LR) interactions, which trigger the activation of downstream signaling pathways. Finally, it affects the activities of transcription factors (TFs) and their target gene expression [11-13]. CCIs mediated by LR interactions have been the most common scenario for the computational study of CCIs in recent years [9].

With the recent advancement of single-cell technologies [14], a large amount of single-cell RNA-sequencing (scRNA-Seq) data has become publicly available [15-21]. scRNA-Seq has enabled new and potentially unexpected biological discoveries compared to traditional profiling methods. These include revealing complex and rare cell populations, uncovering regulatory relationships between genes, and tracking the trajectories of distinct cell lineages in development [19, 20, 22]. Compared to the traditional assays that directly measure protein-protein interactions (PPI), the transcriptomic measurements at single-cell resolution greatly facilitate the computational study of CCIs [23]. With the LR interaction information, scRNA-Seq has demonstrated its effectiveness in exploring CCIs at single-cell resolution.

Many computational tools have been developed to identify CCIs through scRNA-Seq data integration under specific cellular and physiological conditions [24]. These CCI prediction tools, in general, follow a common pipeline, including cell-type classification, LR interaction inference, CCI network construction and CCI visualization. However, each tool has its specific emphasis and algorithmic details. Existing comparative studies of CCI tools mainly report their advantages and disadvantages based on the theoretical analysis [24]. There is a lack of benchmark assessments to understand the performance and effectiveness of the most recent CCI inference tools in real application scenarios. In this work, we attempt to compare and summarize the usage of the recent nine LR-based CCI inference tools using public scRNA-Seq datasets [5, 8, 12, 24-29]. In the following, we first provide an overview of these tools. We next describe the LR resources used by different tools and the benchmark LR interaction database used for the later tool evaluation. We then introduce the three benchmark scRNA-Seq datasets involving transcriptional profiling of 95,145 single cells. Finally, we discuss the systematic evaluation of nine tools in LR interaction prediction and CCI network construction.

### Computational tools for LR-based CCI inference

We compiled nine tools that can predict LR-based CCIs based on scRNA-Seq gene expression measurements, including CellPhoneDB, iTALK, NATMI, PyMINEr, NicheNet, SingleCellSignalR, CellChat, ICELLNET and scMLnet [5, 8, 12, 24-29]. These tools were published in the last four years and run smoothly (Table 1). All these nine tools were developed using either R or python. Almost all the tools except PyMINEr require a curated LR interaction database in addition to raw or normalized Unique Molecular Identifiers (UMIs) counts as input. UMI is a type of molecular barcoding that provides error correction and increased accuracy during sequencing. Among these tools, CellPhoneDB, iTALK, NATMI, PyMINEr, and scMLnet also need the cell type annotation as input. Others perform cell-type annotation by embedding certain cell clustering procedures such as Seurat or K-means in their pipelines and then assuming cluster-corresponding cell types [30]. All of these tools output the predicted LR interaction pairs between cell types. Such LR pairs can then be used to construct CCI networks suggesting the potential communication between cells. In addition, all of them can provide visualization of CCIs. The nine tools are briefly described as follows.

**Table1.** Overview of the CCI inference tools.

| Tools | Reference | Database<br>(# LR<br>pairs) | code | CCI inference | Input: gene<br>expression<br>(normalized<br>/raw data) | Input:<br>Predefine<br>d cell<br>types | Visualization |
| --- | --- | --- | --- | --- | --- | --- | --- |
| iTALK | Wang et al.(2019) | 2,648 | R | CCIs are inferred by finding significant LR interactions based on differentially expressed ligand and receptor identification. | Raw | YES | Circos plots and boxplots of CCI networks |
| PyMINer | Tyler et al. (2019) | NA | Python | CCIs are inferred by identifying autocrine and paracrine signaling through LR-involved pathway analysis. | Normalized | YES | network graph and circular plot of CCIs |
| NicheNet | Browaeys et al. (2020) | 12,652 | R | CCIs are inferred based on the influence of a ligand in one cell type on target genes of a signaling path in another cell type through prior ligand-to-target signaling pathway integration. | Raw | NO | Circos plot of CCIs |
| CellPhoneDB | Efremova et al.(2020) | 1,396 | Python | CCIs are based on identifying statistically significant LR interactions based on their gene expression in involved cell states. | Normalized | YES | Dot plot of cluster combinations |
| NATMI | Hou et al.(2020) | 2293 | Python | CCIs are based on the summation of LR interactions weighted by multiple metrics considering the ligand and receptor expression levels in the two involved cell clusters. | Raw | YES | Heatmap-view, network graph and circular plot of CCIs |
| SingleCellSignalR | Cabello-Aguilar et al. (2020) | 3,251 | R | CCI probabilities are calculated through a nonlinear function of the product of ligand and receptor expressions. | Normalized | No | network graph and circular plot of CCIs |
| CellChat | Jin et al.(2021) | 2,021 | R | CCI probabilities are derived using the law of mass action according to the ligand- and receptor- expression levels averaged over their associated cell groups. | Normalized | No | Alluvial and Circos plots of communication pathways |
| ICELLNET | Noël et al. (2021) | 752 | R | CCI scores are calculated based on the mean expression profiles of their respective ligands and receptors in two cell clusters. | Raw | NO | network visualization of CCIs |
| scMLnet | Chenf et al.(2021) | 2,557 | R | CCIs are inferred by the multilayer network constructed through integrating expression-based reconstruction of LR interaction, receptor-TF, TF-target gene networks. | Raw | YES | Multilayer network visualization of ligand-receptor-TF-target gene links for each cell cluster |

### iTALK

iTALK [25] was developed as a tool for identifying and illustrating CCIs. iTALK was motivated by understanding the crosstalk between tumor cells and other cells in a tumor microenvironment. iTALK manually curated a database that contains 2,648 unique LR pairs. According to the functions of the ligands, iTALK classifies the interacting pairs into four categories, including cytokine/chemokine, immune checkpoint, growth factor and others. With the cell-gene expression matrix and predefined cell types as input, iTALK identifies CCIs as differentially expressed LR pairs. The identification of differential expression is made by integrating existing tools such as DESeq, scde and Monocle [31-33]. iTALK provides various ways to visualize the CCIs, including network, circular and errorbar plots.

### PyMINEr

PyMINEr [28] is an open-source program that can perform multi-facet analysis of gene expression data such as clustering for cell-type identification, differential expression identification, and pathway analyses. Its LR-based CCI inference was illustrated through the prediction of autocrine and paracrine signaling networks. PyMINEr first identifies differentially expressed genes between cell types as cell type enriched genes. Two separate lists of cell-type enriched genes that have the potential to function as ligands and receptors in cell communication are then generated according to their Gene Ontology (GO) annotation and subcellular localization analysis. An LR pair can be defined if the cell type enriched genes are in the two lists and can have physical interaction at the protein level according to the PPI information in StringDB. Such obtained LR pairs are then integrated into the pathway analysis to identify autocrine and paracrine signaling between all cell types. PyMINEr was applied to human pancreatic islet scRNA-Seq datasets, and it was able to identify the BMP-WNT signaling responsible for cystic fibrosis pancreatic acinar cell loss. PyMINEr output an HTML webpage to display all the analysis results, including network graph and circular plot of CCI networks.

### NicheNet

NicheNet [12] attempts to elucidate the functional understanding of CCIs by inferring the functional effect of ligands in the sender cells on the expression of genes in the receiver cells. To do that, NicheNet integrates LR interaction, signal transduction and gene regulatory interaction information. The LR interactions were compiled from KEGG, Reactome, the IUPHAR/BPS Guide to pharmacology, and PPI databases [10, 34-36]. The signaling and PPI information was obtained from multiple pathways and PPI databases such as ConsensusPathDB, Omnipath, and PathwayCommons [37-45]. The gene regulatory interactions were compiled from multiple resources, including TRANSFAC, JASPAR, MSigDB, and so on [46-58]. Individual interactions were organized as weighted networks where LR and signaling networks were combined as a weighted ligand-signaling network, and gene regulatory interactions were converted to a weighted gene regulatory network. A weighted sum of the individual networks was then performed to integrate different data sources. Network operations such as PageRank and matrix multiplication were applied to this integrated network to derive a prior model of ligand-target regulatory potential. After combining the expression profiles of interacting cells, NicheNet can prioritize the regulatory potential of ligands on the target genes. When applied to the HNSCC tumor single-cell data, NicheNet identified TGFB3 as the most probable ligand that regulates the epithelial-to-mesenchymal transition program in a group of malignant cells. The signaling paths from TGFB3 to its target genes were also inferred. NicheNet, when applied to study the CCIs among immune cells using the mouse immune cells, identified antiviral-relevant ligands such as Il27, Ifng and Il12a.

### SingleCellSignalR

SingleCellSignalR [27] curated a LR interaction database, LRdb, from multiple resources such as FANTOM5 [59], HPRD [60], Reactome [61] and HPMR [62]. LRdb requires a potential LR pair with its respective GO annotations being ligand or receptor and involved in the Reactome Pathway database [63, 64]. LRdb contains 3,251 reliable LR pairs. The LR interactions are determined by a regularized product score that is majorly derived from the product of the expression levels of the associated ligand and receptor. A scoring threshold is estimated using multiple benchmark datasets, including a metastatic melanoma dataset, two peripheral blood monoclonal cell datasets, a pan T-cell dataset, a head and neck squamous cell carcinoma dataset. The usage of SingleCellSignalR was demonstrated by the identified LR interactions between cells in a mouse interfollicular epidermis dataset.

### CellPhoneDB

Unlike the previous studies such as iTALK, singleCellSignalR and NicheNet, CellPhoneDB v2.0 considers multiple subunit architecture for both ligands and receptors [5]. CellPhoneDB also curated a LR database, which contains multiple LR subunits. Significant LR interaction pairs are defined based on the likelihood estimation of their respective cell type enrichment. The number of significant LR interaction pairs is used to prioritize the CCIs specific to two given cell types. The CCI-based networks can be constructed to assess cellular crosstalk between different cell types. The cell subsampling method is used to reduce memory usage and runtime.

### NATMI

NATMI (Network Analysis Toolkit for Multicellular Interactions) [29] compiled coonectomeDB2020 that contains 2,293 manually curated LR pairs. NATMI considers a ligand or receptor as expressed if it is expressed in at least 20% of the cells of a given cell type. NATMI then defines an edge weight corresponding to a LR pair using three metrics differing in summarizing the expression levels of a ligand or receptor in cells of the same cell type. For example, one metric is the mean-expression weight, which is defined as the product of the ligand’s mean expression and the receptor’s mean expression in the cells of the cell type under consideration. Based on the edge weights between LR pairs, NATMI constructs a cell-connectivity-summary-network that summarizes CCIs. The criteria can be specified by users, for example, simply counting the number of LR pairs whose weights pass user-defined thresholds. NATMI was applied to the Tabula Muris atlas that contains single cells from 20 organs in mice [27]. Autocrine, intra-organ and inter-organ signaling was identified among 117 cell types in the Tabula Muris dataset. Cellular communities and differential networks were also predicted. NATMI provides a function to visualize the extracted LR pairs and their edge weights.

### CellChat

Like CellPhoneDB, CellChat [8] also incorporates multiple ligand/receptor subunits. It further extends their consideration to additional cofactors such as soluble agonists, antagonists and stimulatory and inhibitory membrane-bound co-receptors. Based on manual curation, CellChatDB was constructed with over 2K LR interactions, each of which is associated with a literature-supported signaling pathway. CCI prediction between a given pair of cell groups is based on a probability calculated by integrating the PPI network and the differential expressed ligands/receptors in the involved cell types. Based on the predicted CCIs, CellChat also performs network analysis on the intercellular CCI networks to identify the dominant roles of different cell types and CCI patterns. CellChat demonstrated its functions using several scRNA-Seq datasets, e.g., the single-cell mouse skin datasets covering embryonic development and adult wound healing stages. With the adult wound healing data, CellChat identified TGF\beta signaling from myeloid cells to fibroblasts, consistent with myeloid cells’ role in literature. CellChat’s pattern recognition module also revealed connections between cells and signaling pathways. For instance, multiple pathways such as ncWNT, SPP1, MK and PROS were identified corresponding to the outgoing signaling of fibroblast cells. With the embryonic day D14.5 mouse skin dataset, CellChat showed its ability to identify CCIs in continuous cell states inferred by the pseudotemporal trajectory.

### scMLNet

Similar to NicheNet, scMLNet [26] integrates multiple types of information, including LR interactions, signaling and gene regulatory interactions as subnetworks to study CCI. scMLNet focuses more on the context-dependent integration by incorporating scRNA-Seq expression data into each subnetwork construction. The constructed subnetworks are output as a multilayer network representing CCIs. scMLNet was applied to a single cell dataset of bronchoalveolar lavage fluid samples in 9 COVID patients and four healthy controls. The CCIs between secretory cells and other cell types enabled the identification of ACE2 regulatory pathways such as PI3K- Akt, JAK-STAT, TNF and MAPK signaling pathways, which were further validated using bulk gene expression data and additional experiments.

### ICELLNET

ICELLNET [24] is a computational framework that can infer CCIs from bulk transcriptomic and scRNA-Seq data. ICELLNET integrated hundreds of literature-annotated and experimentally validated LR interactions as a database. Similar to previous methods, ICELLNET scores the LR interactions using the expression levels of both ligands and receptors in the corresponding cells. One unique feature of ICELLNET is its ability to incorporate gene expression profiles of other cell types not from the same dataset. For example, a cell type can be from the Human Primary Cell Atlas. Briefly, given the expression profiling of a specific cell type, called “central cell”, and that of other cell types provided by the user, called “partner cells”, ICELLNET can predict the potential CCIs between the central cell and partner cells. Due to the incompleteness, complexity and possibilities to cause false CCI predictions, ICELLNET did not integrate pathway and gene regulatory information [65-68]. ICELLNET was applied to human breast cancer-associated fibroblast cells (CAFs), and demonstrated its capability to reconstruct the CCIs and identify the LR interactions between CAFs and 14 other cell types involved in the tumor microenvironment. ICELLNET also provides a few visualization tools for result interpretation.

Even though all the tools can predict CCI networks, they have different considerations in defining LR interactions. We can thus further classify the tools into two categories: pathway-involved and non-pathway-involved. Tools like NicheNet, PyMINEr and scMLnet fall into the pathway-involved category, while others are majorly non-pathway-involved. The pathway-involved tools are primarily motivated by the essential role of intracellular signaling and transcriptional regulation events during CCIs [12, 26, 69, 70]. Therefore, these tools take into account the signaling pathway components by performing pathway integration and analysis when inferring LR interactions and CCIs. Because the current understanding of pathway and gene regulatory mechanism is not complete, the integration of downstream signaling pathways and targets can lead to false predictions [24, 67, 71, 72]. On the other hand, non-pathway-involved tools focus on the transcriptional abundance of the ligands and receptors. Some non- pathways-involved tools such as CellChat and SingleCellSignalR also perform pathway analysis, but only after the CCI inference [8, 27]. Meanwhile, most of these tools utilize the expression levels of ligands and receptors to score the potential LR interactions, which is common in CCI literature [73-77]. Nevertheless, the formulas used to compute the score vary. For example, NATMI offers three different metrics for calculating the LR interaction scores, SingleCellSignalR uses a regularized product, and CellPhoneDB applies modified mean p-value. The corresponding score cutoffs are also defined differently. Although all the tools can generate predictions for important LR interaction pairs, they do not always directly output the LR interaction scores, which is particularly true for the pathway-involved tools.

## Testing data compilation and tool comparison methods

### LR interaction data used for comparison

To evaluate the predicted LR interactions from the nine tools, we downloaded the compiled LR interactions in 23 resources (15 human and eight mice) from recent studies [8, 10, 12, 24, 25, 27, 29, 78-90] (Supplementary Table 1). The majority of the LR interactions were inferred from protein interaction, gene function and pathway annotations in the literature. We then converted the LR interaction from human and mouse to be the same unified human gene symbol using the bioMart package [91]. After the conversion, the number of LR interactions in the 23 resources varied from 456 to 49,323. Although many LR interactions are shared among these resources, much more LR interactions are unique to individual resources. To find the most reliable LR interactions, we calculated the overlap of the LR pairs between resources (Fig. 1). We defined the most reliable LR interactions as those occurring in at least four resources. This definition resulted in 3,779 LR interaction pairs forming the consensus LR interaction database (C-LRI). We used the C-LRI database as a benchmark dataset to evaluate the LR predictions from different tools (Supplementary File S1).

**Figure 1.**
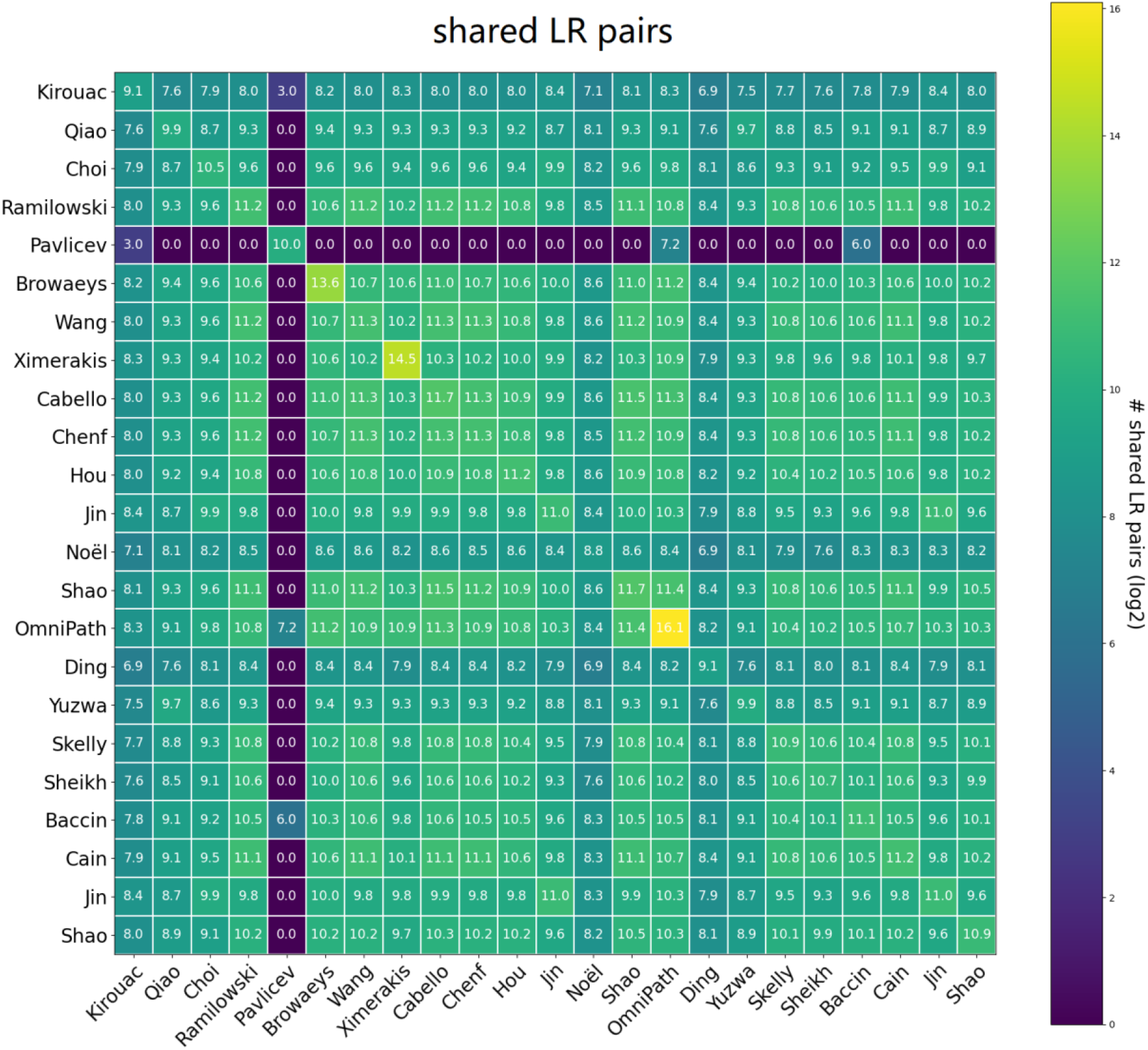
The overlap of the LR pairs in the 23 resources

#### scRNA-Seq datasets

We evaluated the nine tools using three scRNA-Seq datasets (Table 2), involving 15 scRNA-Seq samples.

**Table 2.**
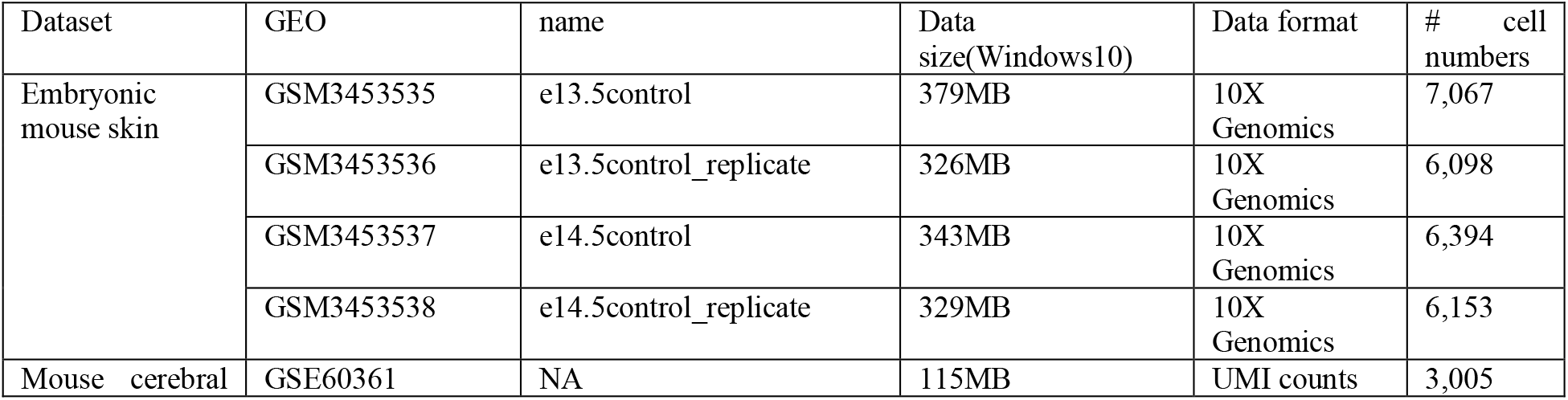
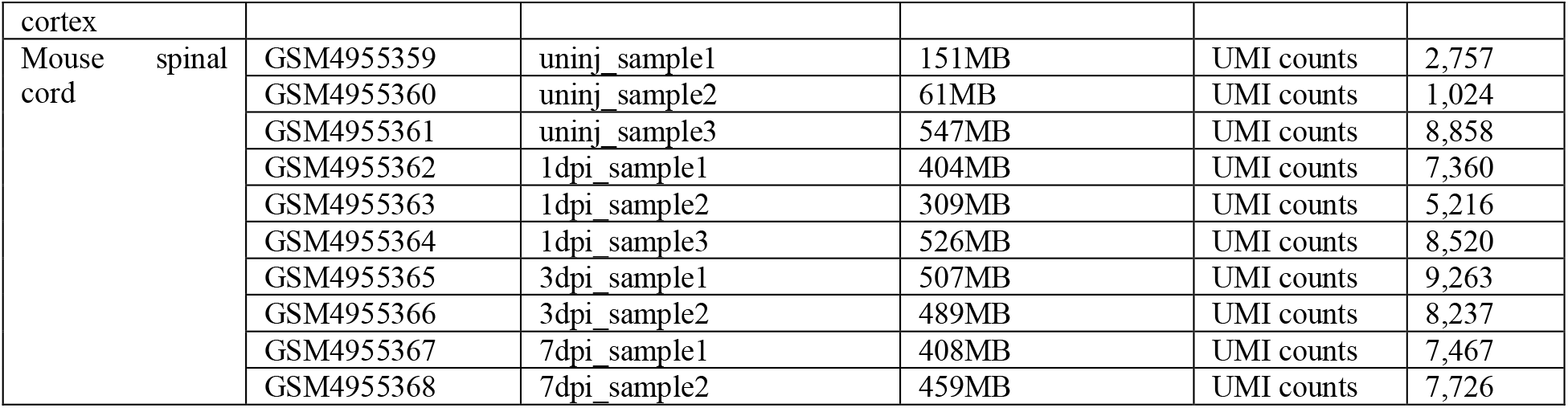
Details of the 15 samples

The first dataset is the scRNA-Seq data from embryonic mouse skin [92]. This scRNA-Seq data corresponds to the gene expression measurement of single cells of embryonic dorsolateral/flank skin at embryonic days 13.5 and 14.5. We obtained two biological replicates for both days (GSM3453535, GSM3453536, GSM3453537, GSM3453538). We then performed the cell-level filtering by removing the cells with UMI counts less than 2500 or greater than 50000 [8]. We also removed cells with their gene number less than 1000 and the fraction of mitochondrial counts greater than 20%.

The second dataset is the mouse cerebral cortex scRNA-Seq data [93], which measures gene expression in 3,005 high-quality single cells isolated from the mouse cerebral cortex (GSE60361). It contains the main cell types in the hippocampus and somatosensory cortex. We then filtered out unreliable genes based on the total number of reads per gene and only kept the genes detected in more than 30 cells [94].

We also compared the nine tools with the third scRNA-Seq dataset from the mouse spinal cord (GSE162610) [95]. This dataset measures gene expression in all cell types of the uninjured and injured spinal cord of wild-type mice at one day, three days, and seven days after injury. The data contains 10 samples including three uninjured samples, three 1dpi, two 3dpi and two 7dpi samples (GSM4955359, GSM4955360, GSM4955361, GSM4955362, GSM4955363, GSM4955364, GSM4955365, GSM4955366, GSM4955367, GSM4955368). We used the processed dataset by the original paper.

#### Compare predicted LR pairs

For a given scRNA-Seq sample, the LR interactions between specific cell types were compared. Cell clusters were first obtained by Seurat [30]. Each cell cluster was then annotated to a cell type by cross-checking the cells’ annotation in the original literature reference. Each CCI tool was run to output the top LR predictions between cell type-annotated clusters. Given such obtained LR interactions between any two cell types, all the tools were compared regarding the predicted number of LR interactions. The predicted LR interaction pairs were also compared against the C-LRI database. The LR pairs were used to infer CCIs for each tool. These CCIs were then compared across different tools. To determine how cell clustering results affected the CCI inference results, we enumerated the resolution parameter of Seurat. Note that the resolution parameter is associated with the Seurat clusters’ granularity, where a larger resolution value corresponds to a larger number of clusters.

#### Compare the inferred CCI networks

For each CCI inference tool, a CCI network was constructed based on the predicted LR interactions. In a CCI network, each node represents a cell type, and each edge corresponds to the LR-based CCIs between two cell types. Similar to ICELLNET [24], we defined the pairwise dissimilarities *d*^α,β^ between two CCI networks α and β as:

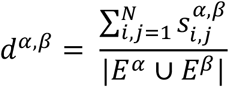

With

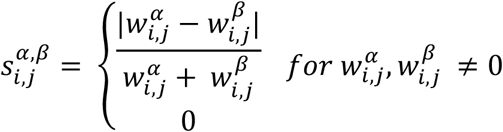

where 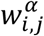 is the weight of the directed edge from nodes *i* to *j* in the CCI network α, *N* is the total number of nodes, *E*^*α*^ is the set of edges in the CCI network α, |*E*^*α*^ ∪ *E*^*β*^| is the cardinality of the union of all edges in CCI networks α and β. Note that, the edge weight can be defined as the number of predicted LRs between two corresponding cell types.

#### Compare the sensitivity by subsampling

To compare the sensitivity of predicted communications to the original input data, we used the ‘geometric sketching’ approach to perform subsampling of the scRNA-Seq datasets [96]. Geometric sketching is a data sampling approach that has been shown effective in using a small portion of the original single-cell data to represent the full data and consequently reduce the data volume. Compared with the commonly used down sampling strategies, the geometric sketching method can keep the rare cell types and maintain the heterogeneity of the original data. To calculate the sensitivity scores for the tools, given a CCI inferred from the sampled data, if it is also inferred from the original data, it is defined as a true positive. Any CCI predictions from the original data but not in the sampled data are false negatives. Such sampling analysis was used to evaluate the consistency between tools rather than the accuracy of each tool.

### Running time and memory usage

We collected the runtime and memory usage of the nine tools on a dedicated machine. All the tools were run on the same Ubuntu 18.04 Long Term Support supporting computer with Intel core i7-10875H CPU @2.3 GHz 16 cores and 128 gigabytes memory. We used the python ‘time’ package to record the time used for each tool. Briefly, we recorded the time between the tool command start and the tool command end. We used the ‘top’ command to monitor the memory used when the tool was running for memory usage. The maximum used memory was selected as the memory usage for each tool. When we ran the code, we tried to suspend all other activities to make sure we could obtain the actual maximum memory usage.

### Comparison of the nine tools using mouse skin and cortex data

#### Comparison of the predicted LR interaction pairs

We found a large inconsistency in the number of predicted LR interactions among the different tools. The number of LR interaction pairs predicted by the nine tools for a given sample often varies from dozens to thousands. To make the predicted LR pairs from different tools comparable, we selected the top predictions accordingly for each tool. Towards this goal, we kept all the LR pairs from CellChat, CellPhoneDB, scMLnet, and SingleCellSignalR, while only selecting the top 10% of LR pairs from iTALK, ICELLNET, NicheNet and NATMI. As the PyMINEr predicted much more than others, we kept the top 1% of its predicted LR pairs.

After selecting the top LR pairs, the difference of the predicted LR numbers is lessened but persists. For example, for the embryonic day 13.5 control replicate sample (GSM3453536) in the Embryonic mouse skin dataset, the number of predicted LR interaction pairs between fibroblast type A (FIB-A) and fibroblast type B (FIB-B) cells still is varied by tools ranging from 12 to 192 (Fig. 2A). This situation is similar for all the four samples in the mouse skin dataset and the mouse cerebral cortex sample. In the mouse cortex sample, the number of LR interactions from the Pyramidal CA1 to oligodendrocytes cells varies from 44 (CellChat) to 235 (ICELLNET) (Fig. 2C). CellChat, scMLnet and SingleCellSignalR predicted fewer than a few dozen LR interactions, while iTALK, ICELLNET and NATMI often output more than one hundred LR interactions based on our selection criteria above. For different tools, not only does the predicted number of LR interactions differ for the same sample but also the number distribution of LR interactions across different samples. For example, the CellChat, CellPhoneDB, PyMINEr predicted most LR pairs from the embryonic day 14.5 control sample (GSM3453537) in the mouse skin dataset, while SingleCellSignalR predicted the least LR interactions in the same sample.

**Figure 2.**
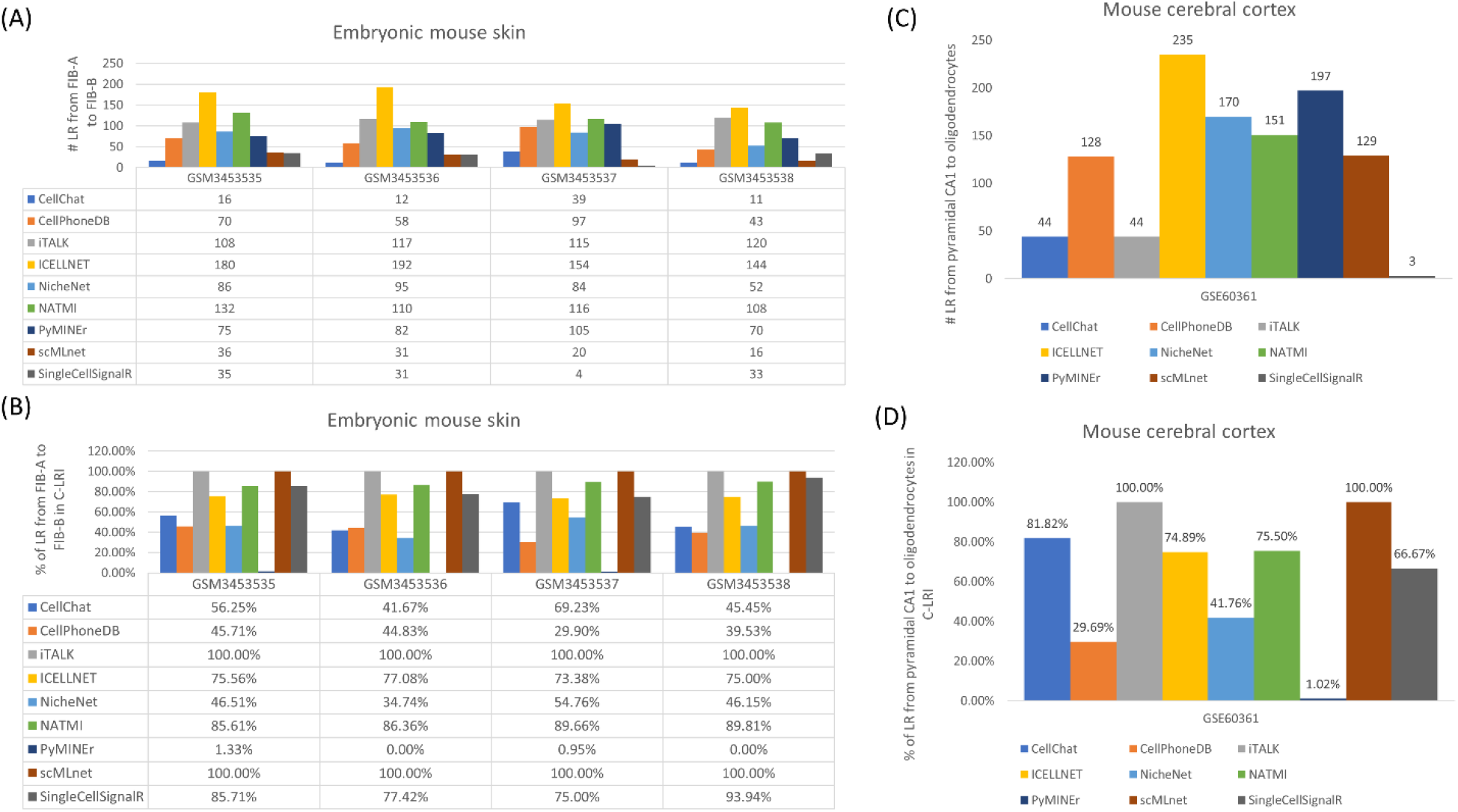
Comparative performance of tools on different datasets. (A) The number of predicted LR pairs from FIB-A to FIB-B cell types in embryonic mouse skin samples. (B) The number of predicted LR pairs in C-LRI from FIB-A to FIB-B cell types in embryonic mouse skin samples. (C) The number of predicted LR pairs from pyramidal CA1 to oligodendrocytes cell types in mouse cerebral cortex. (D) The number of predicted LR pairs in C-LRI from pyramidal CA1 to oligodendrocytes cell types in mouse cerebral cortex.

A closer examination of the LR pairs shows few predictions are shared between tools (Supplementary Figs. 1&2). Although iTALK, iCELLNET and NATMI all had over one hundred pairs of LR interaction predicted, they only share 20-30% of their predictions. For example, for the embryonic day 13.5 control replicate sample (GSM3453536), iTALK shares with ICELLNET and NATMI 23 (19.7%), 39 (33.3%) of its 117 LR predictions, respectively.

Meanwhile, among its 192 LR predictions, ICELLNET only has 23 (12%), 38 (19.8%) LR pairs in common with iTALK and NATMI, respectively. On the other hand, for the tools that frequently predicted fewer LR pairs in a given sample, the overlap between the predicted LR pairs is even less. For instance, out of the 31 predicted LR pairs, SingleCellSignalR only has one LR pair in common with scMLnet.

We further compared the LR predictions against the C-LRI database (Fig. 2B). We found that all the predictions from iTALK and scMLnet are in the database C-LRI. A large percentage of NATMI, SingleCellSignalR and ICELLNET predictions are in the C-LRI. For example, for the embryonic day 13.5 control replicate sample (GSM3453536), 86.36% of the NATMI’s 110 LR predictions, 77.42% of the SingleCellSignalR’s 31 predictions and 77.08% of the ICELLNET’s 192 predictions were found in the database C-LRI. In contrast, CellChat, NicheNet, CellPhoneDB have a smaller percentage of overlap with the C-LRI data (41.67%, 34.74% and 44.83%, respectively). The same pattern is approximately followed by the mouse cortex sample where iTALK and scMLnet predicted all C-LRI LR pairs, while ICELLNET and NATMI predicted more C-LRI pairs than others taking into account the number of predictions (Fig. 2D). Among the nine tools, PyMINEr is the only exception in that it has little overlap with the C-LRI data. Further inspection shows that PyMINEr’s predictions are largely PPIs that are not often annotated LR interactions.

The above observations were made when the resolution parameter of Seurat was set as 0.4 (corresponding to 20 clusters assigned to seven cell types). We also investigated how cell clustering with various Seurat resolution settings might change the LR interaction prediction of the nine tools. To do that, we set the resolution to be 0.1, 0.3, 0.5, 0.7 and 0.9, corresponding to 8, 15, 20, 22 and 22 clusters, respectively. We then compared all LR pairs predicted by each tool. We found that cell clustering impacted the tools differently. Take the mouse cerebral cortex sample as an example. Although the resulted cluster number changes, the overall number of LR pairs remains similar for most tools (Supplementary Fig. 3). Only ICELLNET and NATMI have little difference in resolution 0.1 compared with other parameters. However, when examining LR interactions between specific cell types, we observed that the number of LR pairs consistently increases with the higher resolution setting, except SingleCellSignalR. For example, we obtained LR interactions between cell types pyramidal CA1 and pyramidal SS for each tool under different Seurat resolution parameter settings (Fig. 3). We observed the number of CellChat-predicted LR pairs is 15, 21, 24, 26 and 26 for resolution 0.1, 0.3, 0.5, 0.7 and 0.9, respectively. However, the number of SingleCellSignalR-predicted LR pairs is 15, 21, 24, 26 and 26 for resolution 0.1, 0.3, 0.5, 0.7 and 0.9, respectively. The distributions of LR interactions for the nine tools are similar for other cell types, e.g., the number of LR pairs between cell type pyramidal CA1 and oligodendrocytes (Supplementary Fig. 4). With more clusters generated, the same LR pair can be predicted multiple times by multiple clusters corresponding to the same cell types, leading to more predicted LR pairs. The exception of SingleCellSignalR might be due to its specific procedure to select significantly expressed genes for consideration in LR prediction. Therefore, variations in cell clustering and cell-type classification can impact the prediction of LR interactions.

**Figure 3.**
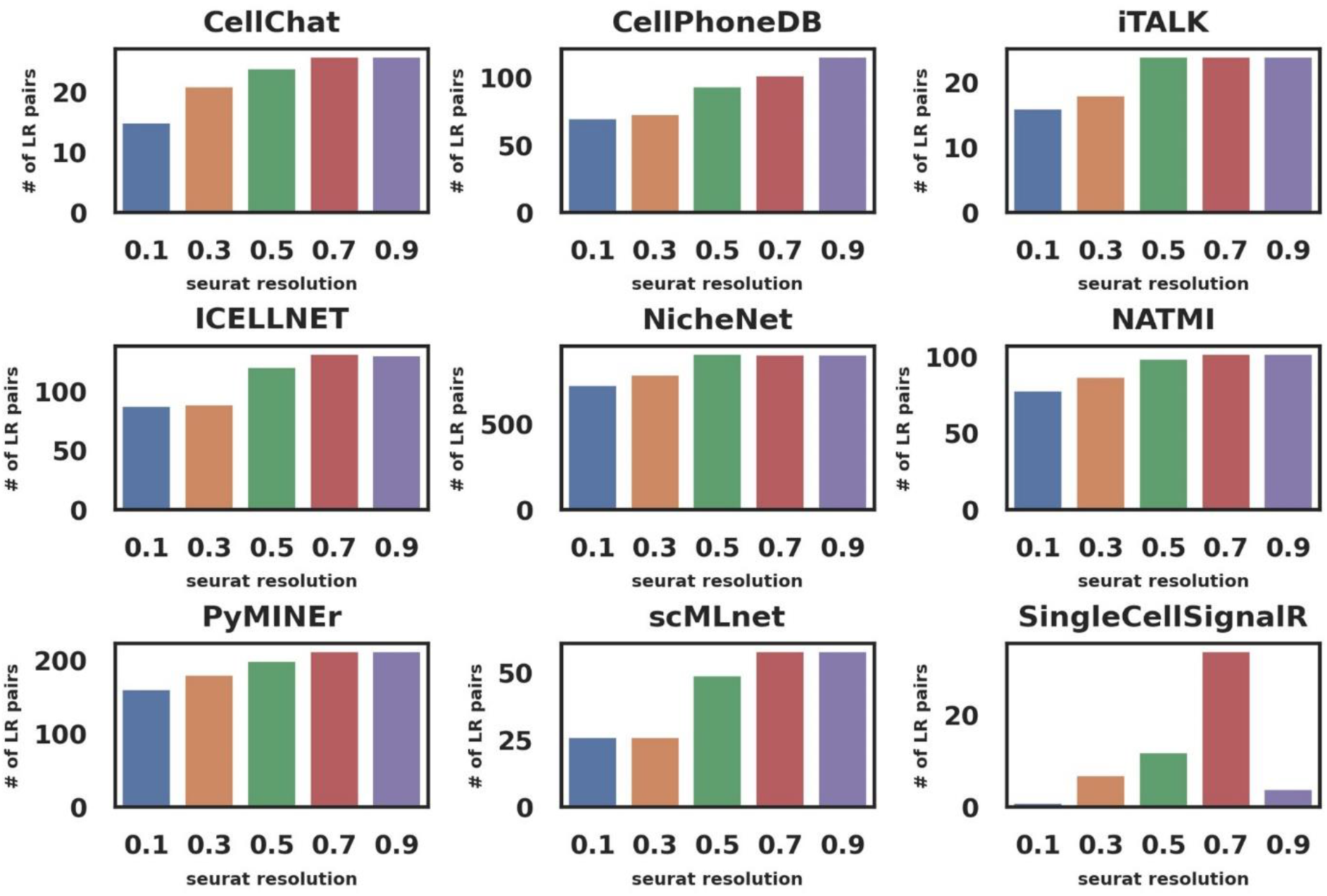
The number of LR pairs between pyramidal CA1 and pyramidal SS cell types in mouse cerebral cortex, predicted by the nine tools under different Seurat resolution parameter settings.

**Figure 4.**
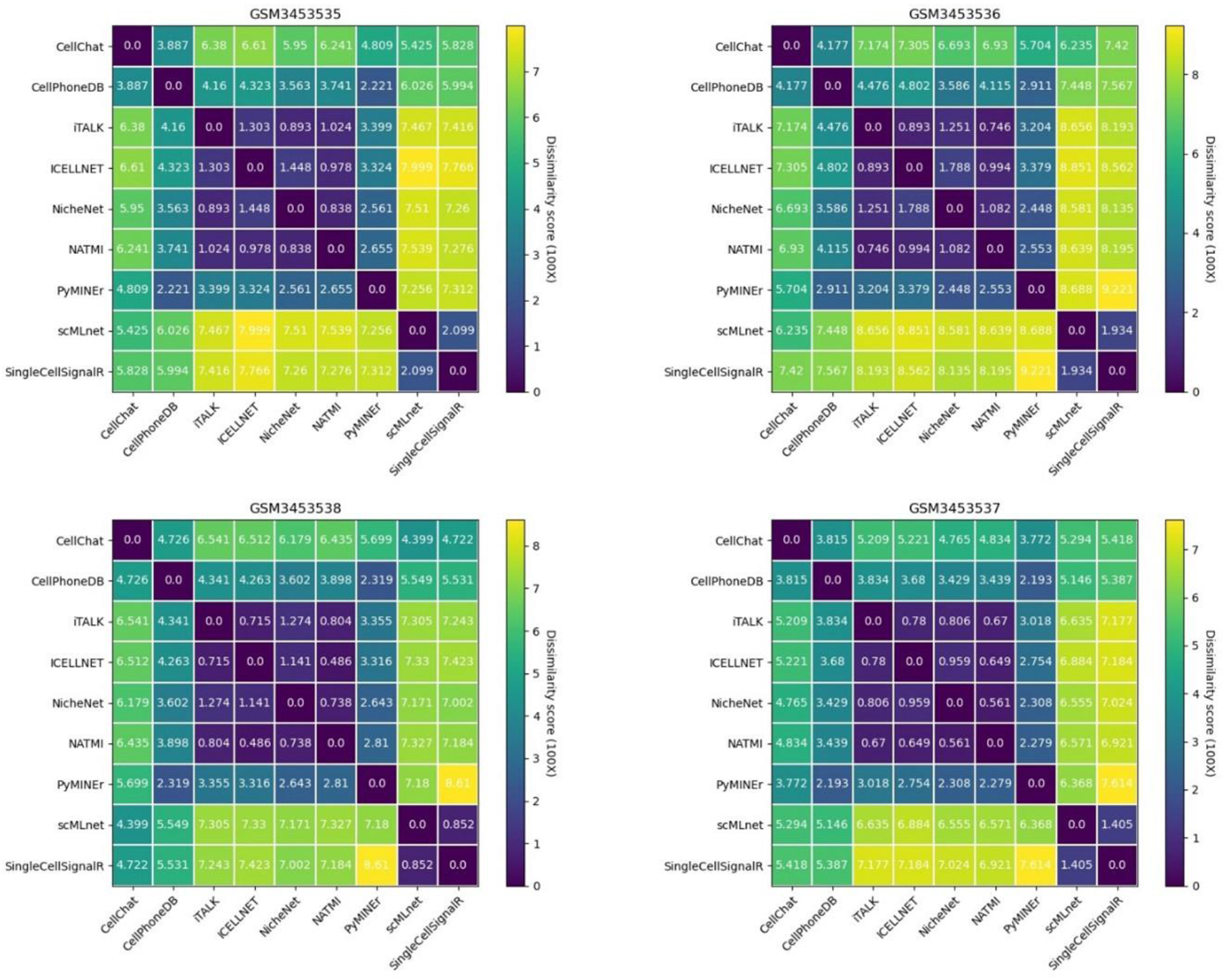
CCI network dissimilarity scores between different tools corresponding to the four samples in the embryonic mouse skin dataset.

#### Comparison of the predicted CCI networks

We compared the CCI networks generated by the nine tools using the dissimilarity score (Section “Testing data compilation and tool comparison methods”). We found that the CCI dissimilarity patterns are conserved between different samples. Fig. 4 shows the dissimilarity scores between tools applied to the four samples in the embryonic mouse skin dataset. We observed that ICELLNET, NATMI, iTALK and NicheNet often have similar CCI outputs, scMLnet and SingleCellSignalR, in general, have similar CCIs. However, CCI networks from SingleCellSignalR are often very different from those produced by other tools. For example, the mean dissimilarity score between PyMINEr and the SingleCellSignalR was 8.12. In contrast, ICELLNET and NATMI have an average score of 0.78. The dissimilarity pattern between tools is followed by the mouse cortex sample as well (Supplementary Fig. 5).

**Figure 5.**
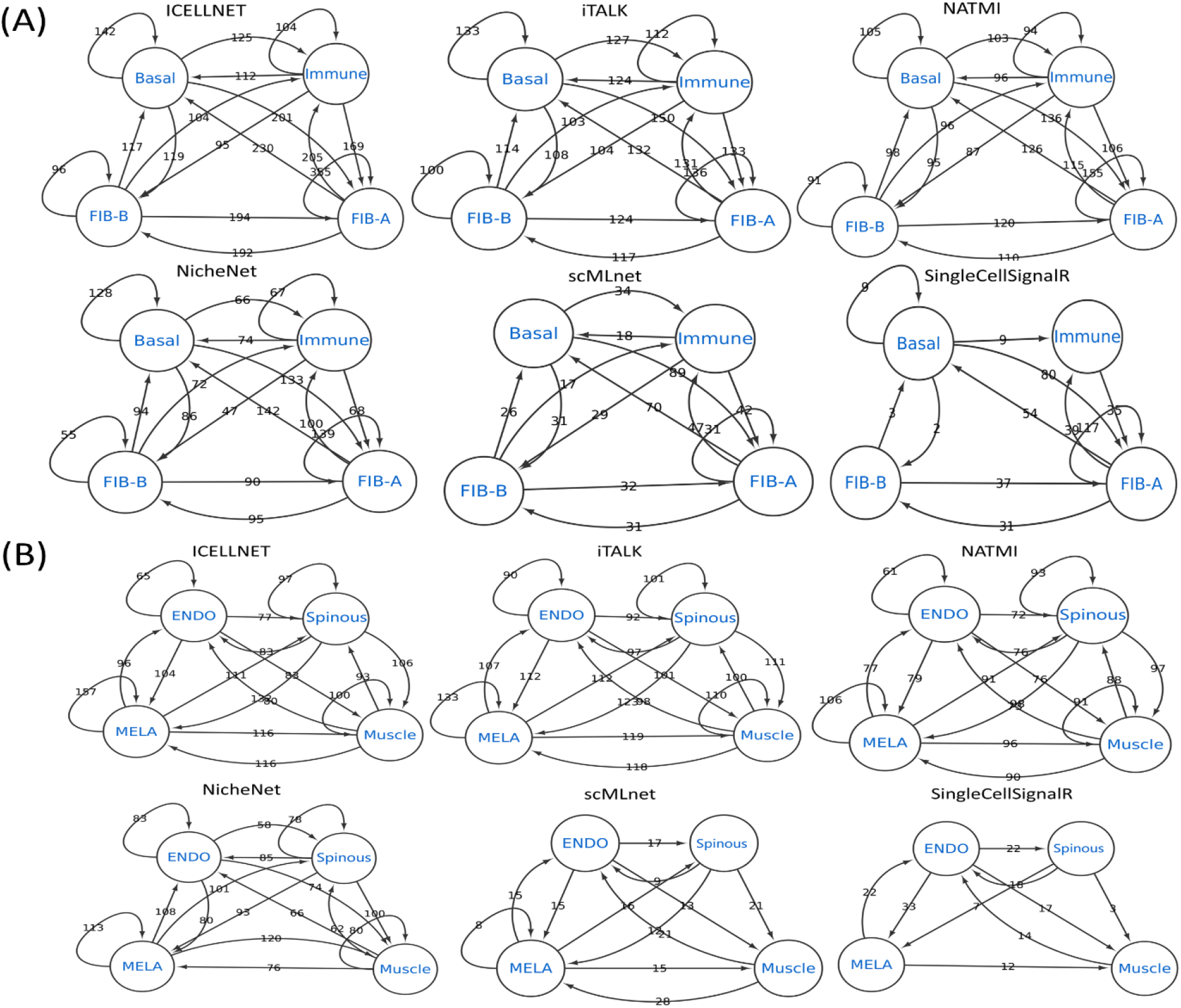
CCI network examples of selected five tools on embryonic mouse skin dataset (GSM3453536). (A) The CCI subnetwork corresponding to the basal, immune, FIB-A and FIB- B cell types. (B) The CCI subnetwork corresponding to the endothelial, spinous, MELA and muscle cell types.

Further inspection of the CCIs between different tools shows that the difference of CCI networks is largely derived from the LR interaction predictions. For the embryonic day 13.5 control replicate sample (GSM3453536) in the mouse skin dataset, we observe the same CCI edges identified by ICELLNET, iTALK, and NATMI with slightly varied edge weights. In contrast, the SingleCellSingalR and scMLnet generally have much smaller edge weights and fewer CCI edges between cells (Fig. 5). In the CCI network corresponding to SingleCellSignalR, the signaling from immune cells to basal cells was not inferred due to the lack of LR interaction predictions. For the same reason, the communication from muscle cells to MELA cells was not inferred by the SingleCellSignalR. The same observation was made for the mouse cortex sample. A larger number of LR predictions, in general, leads to a denser CCI network. Tools such as SingleCellSignalR and scMLnet that tend to predict a smaller number of LRs can result in a lower CCI detection rate. On the other hand, tools such as ICELLNET, iTALK and NATMI tend to predict a larger number of CCIs, which can be associated with lower confidence in the predictions.

We performed the Geometric sketching subsampling procedure to compare the sensitivity of CCI predictions by the nine tools (Section “Testing data compilation and tool comparison methods”). Briefly, we separately sampled 90%, 80% and 70% of the total number of cells in each data. We then ran the nine tools using the sampled data as input. We computed the precision, recall, and F1 scores based on the running results (Supplementary Tables 2&3). The precision is higher with the increasing percentage of subsampling for both datasets. In general, ICELLNET and NATMI have higher subsampling precision and recall overall compared with scMLnet and SingleCellSignalR (Figs. 6A&B).

**Figure 6.**
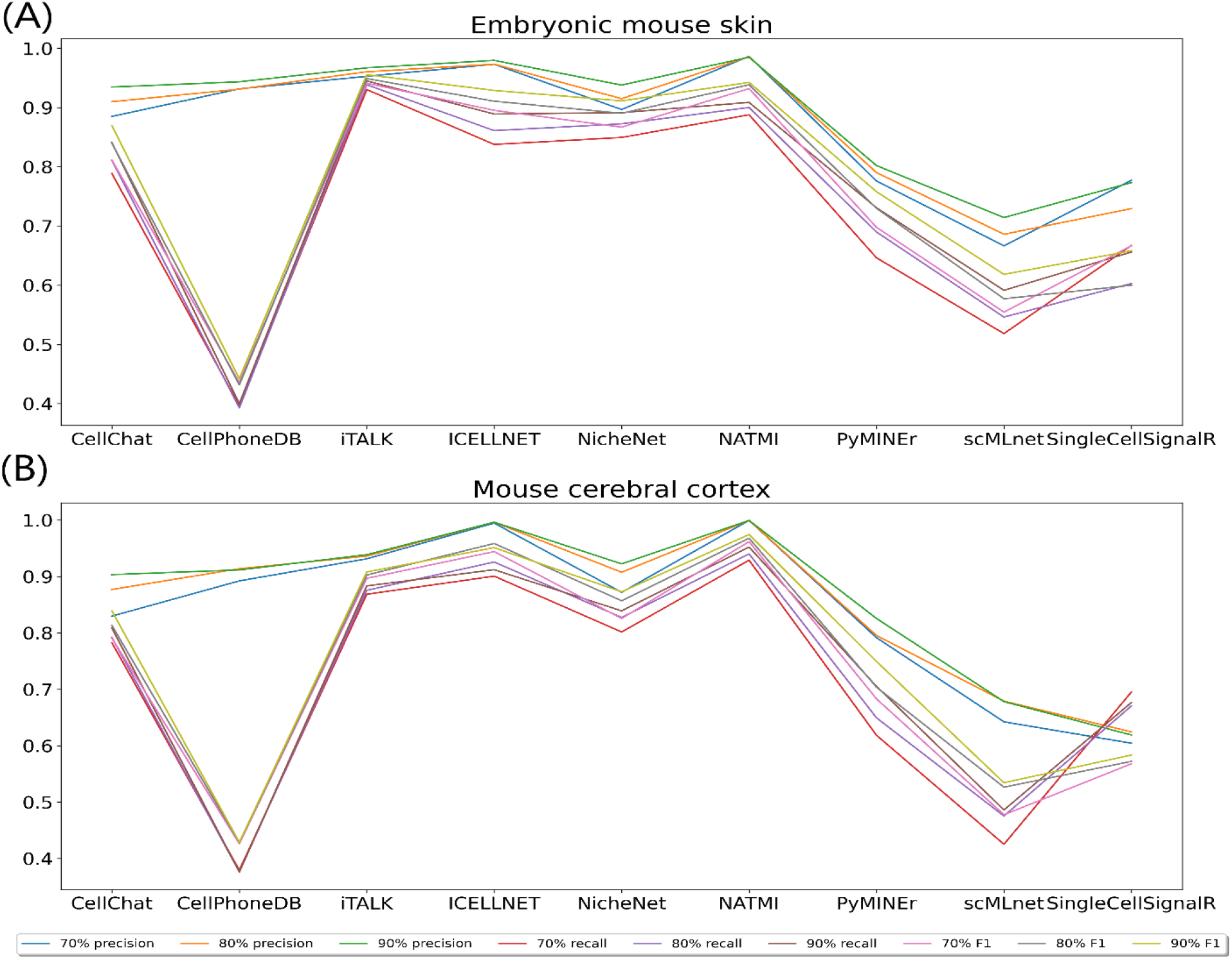
Subsampling sensitivity scores of tools on different datasets. (A) Embryonic mouse skin data. (B) Mouse cerebral cortex data.

### Comparison of the nine tools using mouse spinal cord injury data

We performed the tool comparison on the recently published mouse spinal cord data. The scRNA-Seq data with ten wild-type mice samples correspond to the transcriptional profiling of the uninjured and injured spinal cord at 1 dpi, 3 dpi and 7 dpi. For mouse spine cord samples, we set Seurat parameters to discover the 15 cell types based on the original UMAP plot [95].

Similar to the mouse skin and cortex datasets, the nine tools predicted different numbers of LR interactions. For example, the number of LR interactions from Microglia cells to Endothelial cells varies from 11 (CellChat) to 242 (iTALK) for the “uninjured sample3” (GSM4955361) (Fig. 7A). Consistent with the observations from mouse skin and cortex samples, iTALK, ICELLNET and NATMI often predicted hundreds of LR interaction pairs. The top 10% pairs are counted over one hundred. However, the predicted LR pairs are largely not overlapping. For example, for the “uninjured sample3” (GSM4955361), iTALK shares with ICELLNET and NATMI 31 (12.8%), 50 (20.7%) of its 242 LR predictions, respectively. Meanwhile, ICELLNET only has 31 (17.8%), 46 (26.4%) of its 174 LR predictions in common with iTALK and NATMI, respectively. While NATMI has nearly 40% of its 127 predictions overlapping with iTALK’s 242 predictions, SingleCellSignalR has 74 predictions, with only six overlapping with ICELLNET, five overlapping with NATMI, and zero overlapping with CellChat, CellPhoneDB and PyMINEr. Although CellPhoneDB has 54 predictions, it shared a few with other tools. This situation is similar across all the ten samples (Supplementary Fig. 6). This observation again suggests the large inconsistency of LR interaction ranking and predictions.

**Figure 7.**
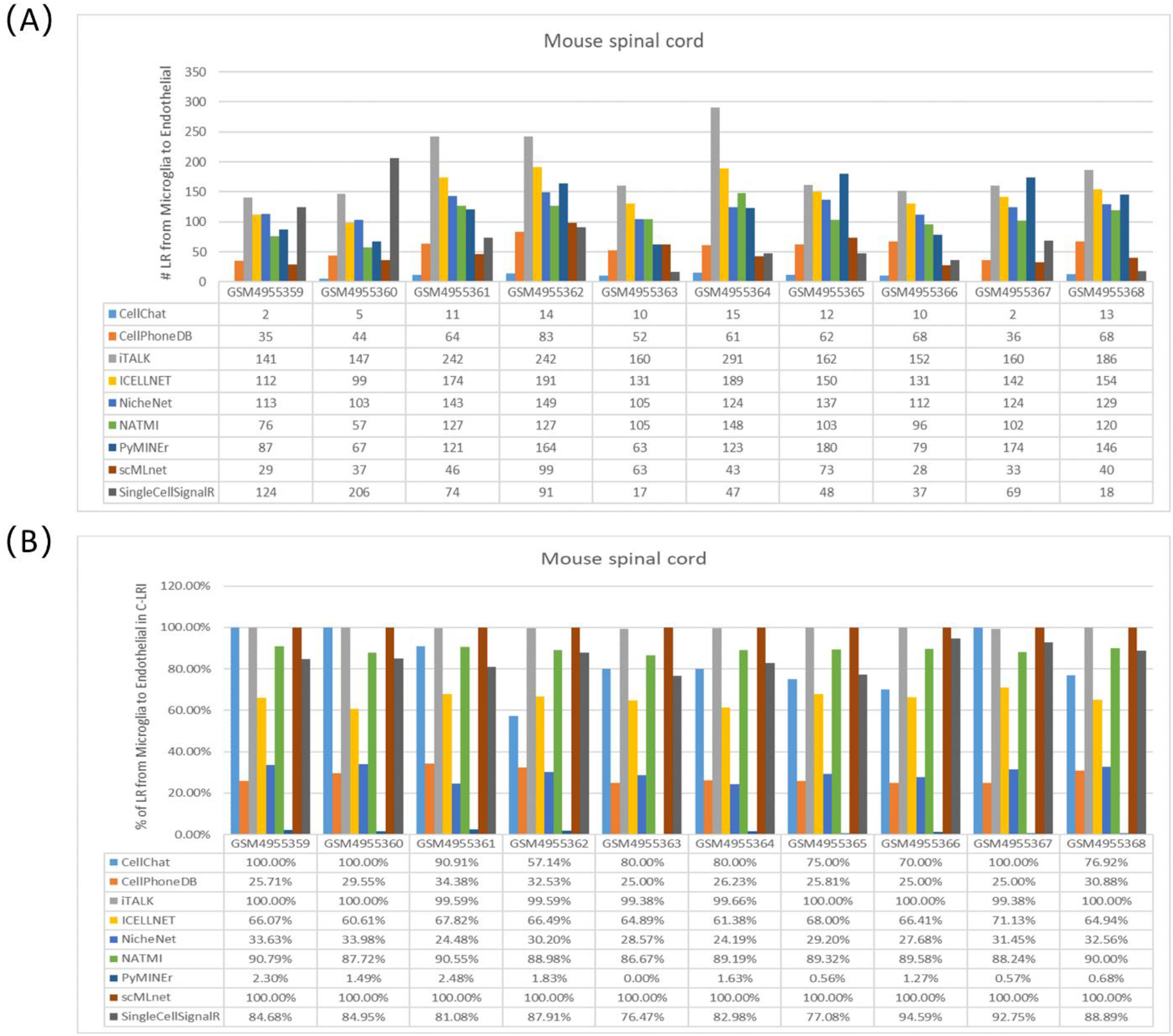
Comparative performance of tools on mouse spine cord injury dataset. (A) The number of predicted LR pairs from microglia to endothelial cell types. (B) The number of predicted LR pairs in C-LRI from microglia to endothelial cell types.

Comparing the nine tools’ LR pair predictions to the C-LRI database for all the samples, we found almost all the predictions from iTALK and scMLnet, the majority of the predictions by CellChat, NATMI, SingleCellSignalR and ICELLNET are in C-LRI. At the same time, NicheNet and PyMINEr have a lower percentage of their predictions in C-LRI (Fig. 7B). Take the “uninjured sample3” (GSM4955361) as an example, 90.91% of the 11 predictions of CellChat, 90.55% of the NATMI’s 127 predictions, 81.08% of the SingleCellSignalR’s 74 predictions and 67.82% of the ICELLNET predictions are also in C-LRI. In contrast, NicheNet, CellPhoneDB and PyMINEr were only found 24.48%, 34.38%, and 2.48% of their 143, 64 and 121 predictions in C-LRI. The resulted statistics for each tool are similar across the other nine samples (Fig. 7B). Considering the total number of LR predictions, NATMI’s LR predictions are most consistent with the C-LRI information, indicating that NATMI is most likely to predict well-annotated LR interactions.

The large variance of LR interaction predictions by different tools has an immediate impact on the study of LR interaction dynamics. We compared the nine tools to see how LR interactions between certain cell types change with the days passed the injury. Multiple tools consistently identified some dynamics of LR interactions. For instance, the THBS1-CD47 interactions between the monocyte and endothelial cells were not found in uninjured mice but were identified in injured mice by ICELLNET, NATMI, PyMINEr and ITALK. It is also interesting that the SPP1-ITGA5 interactions between microglial and endothelial cells were not discovered in the uninjured mice, injured mice at 7 dpi, but only be found in the injured mice at 1 dpi and 3 dpi. This discovery was made consistently by CellChat, PyMINEr, iTALK and scMLnet. Also, the TGFB1-ENG interactions between microglial and endothelial cells became identifiable by ICELLNET, NATMI and iTALK starting at 3 dpi and continuing into 7 dpi. However, most LR pairs were not supported by the majority of the nine tools. For example, the GDNF-GFRA2 interactions between neutrophil and endothelial cells only occur at 1 dpi, identified by ICELLNET and iTALK, but not by other tools. In some cases, the discoveries of the LR dynamics patterns conflict with each other. For example, GNAI2-ADORA1 interactions between neutrophil and endothelial cells were discovered by NATMI at 1 dpi and 7 dpi, but not at 3 dpi and uninjured mice. In contrast, the same LR interaction was identified by SingleCellSignalR only at 3 dpi. In general, we observed that ICELLNET, iTALK and NATMI have high similarities in terms of LR dynamics identification.

Comparison between the CCI networks generated by the nine tools shows CCI dissimilarity patterns are largely conserved between different samples (Supplementary Fig. 7). Again, we observed that ICELLNET, NATMI, iTALK and NicheNet often have similar CCI outputs, while scMLnet and SingleCellSignalR generally have similar CCIs. We also tested the robustness of the nine tools on the mouse spinal cord data with the Geometric sketching approach. We found iTALK and NATMI often show better sensitivity than scMLnet and SingleCellSignalR (Supplementary Table 4).

### Runtime and memory analysis

We compared the runtime and memory cost of the nine tools. We found the running time varies from seconds to hours. For example, using the mouse cerebral cortex dataset as an example, the iTALK used the shortest time (∼20 seconds), while scMLnet took the longest time (8 hours) (Fig. 8). The variation in the time cost is largely attributed to the specific algorithms for CCI inference. For instance, iTALK predicts LR pairs among all cell types simultaneously, while scMLnet and NicheNet consider LR pairs for two cell types one at a time. In certain cases, such as PyMINEr and SingleCellSignalR, additional outputs such as networks and images can make the process longer. The running time is also affected by the number of identified cell clusters.

**Figure 8.**
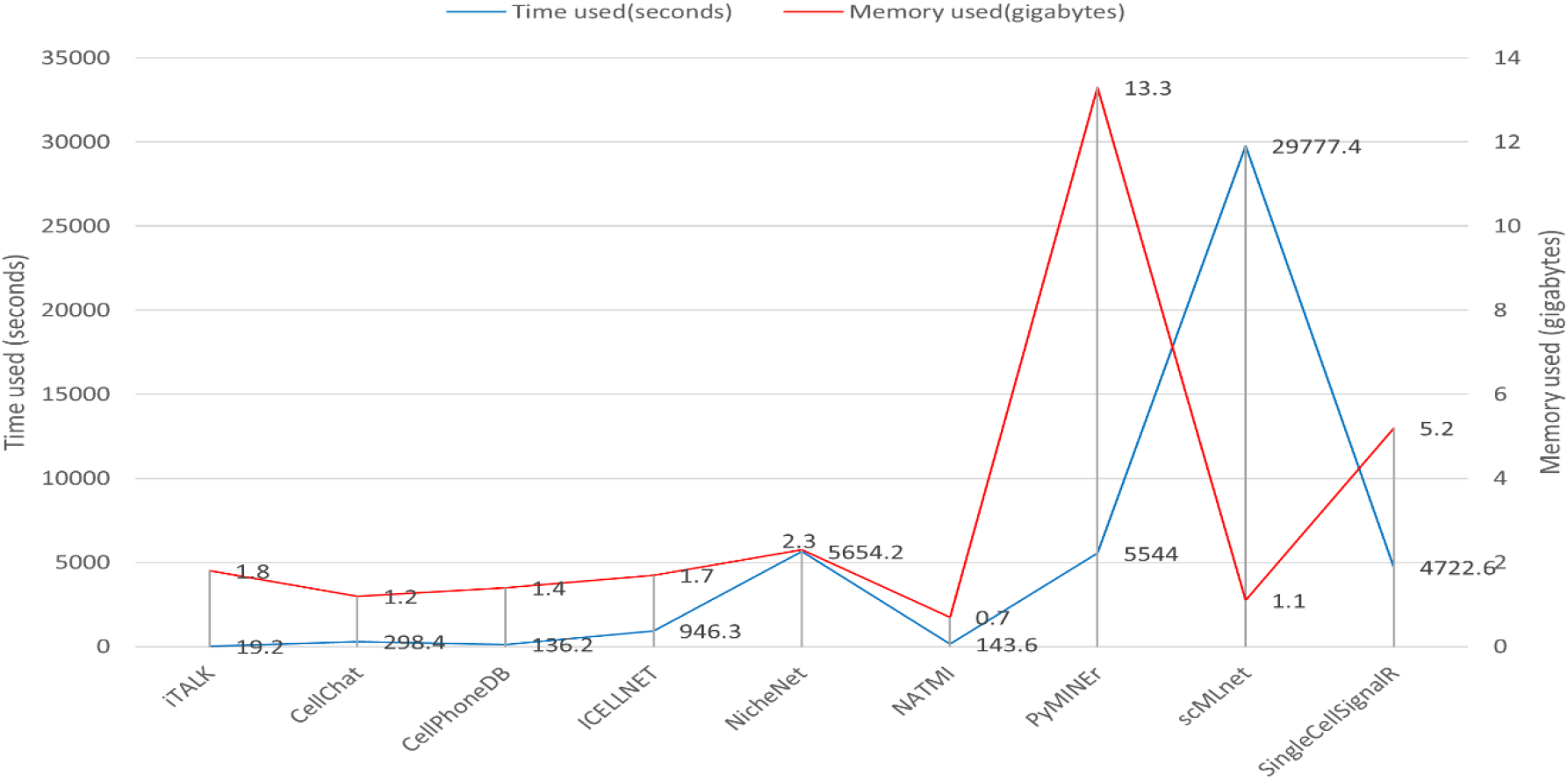
Running time and memory usage.

We also estimated the memory usage of each tool. Almost all of the tools’ memory usage is between 0.5 to 2.5 gigabytes (GB) except for PyMINEr and SingleCellSignalR, which cost 13.3 GB and 5.2 GB, respectively. For PyMINEr, the calculating of correlation between gene pairs can be memory-intensive. For SingleCellSignalR, a large memory is required to obtain a signature cell matrix by computing the average gene signature expression across all the cells and signatures.

### Conclusions and outlook

We studied the nine most recent LR-based CCI prediction tools to understand cell communication at a single-cell level. We provided a side-by-side comparison scenario among the nine tools regarding LR interaction resources, required input, LR output, CCI inference, run time, memory consumption and visualization capability.

The number of predicted LR interaction pairs can significantly differ among the nine tools in terms of LR interaction predictions. Compared with the other tools, which often predict hundreds of LR interactions, CellChat and SingleCellSignalR generate dozens of LR interaction predictions for the testing datasets. We found that iTALK, ICELLNET, NicheNet and NATMI provide a relatively more consistent number of LR interaction predictions with each other than others. However, their predicted LR interaction pairs often do not form consensuses. In terms of the CCI inference, it is directly impacted by the previous LR interaction prediction results. Naturally, we observed that tools with similar LR predictions result in similar CCI networks. We tested the robustness of these tools by performing subsampling of the original input data. We found that most of the current tools have good robustness regarding data selection.

From the above analysis of the 15 samples containing nearly 100K cells, we observed a large discrepancy among the nine tools in LR predictions and the subsequent CCI inference. One reason can be that different tools run on their LR databases, and most of the tools do not allow user-specified LR databases in their current pipeline. We constructed a more reliable LR interaction database C-LRI containing LR interaction pairs supported by at least four literature resources to compare the predicted LR interactions. Therefore, LR pairs in C-LRI are mostly well-annotated LR interactions. We used C-LRI as one way to benchmark the LR predictions from these tools. This benchmark data provided us with additional insight into the tools. For instance, PyMINEr predicted a decent number of LR interaction pairs for our test data, but few of PyMINEr’s predictions are in the C-LRI database. We need to note that even for LR predictions not in C-LRI, one can not claim them as false predictions. However, it does provide an intuition of the LR prediction accuracy, assuming well-known and relatively-new LR interactions are equally distributed.

We tested the sensitivity of LR predictions by enumerating cell clustering/classification at different resolutions. We found that most of the tools generated more LR predictions when more clusters were obtained. One possible explanation is that as the number of clusters increases, the number of possible cell-type pairs increases, and the number of predicted LR interactions also increases. This could indicate issues relevant to multiple hypothesis testing. Meanwhile, because clusters often correspond to cell type classification, the clustering results would affect the CCI inference in the later stage. Besides choosing clustering algorithms that are robust to parameter settings, knowing the approximate number of cell types in the data would increase the confidence of the inferred CCIs. However, accurate specification of cell-type composition for a given sample is often unavailable and requires domain expertise.

We found that the number of single cells in a given sample also affects the number of identified LR pairs for tools, such as iTALK, for which fewer cells often corresponds to fewer LR predictions. This is also evident when comparing the mouse skin samples and the mouse cortex sample, where the mouse cortex sample contains a smaller number (3,005) of high-quality single cells. We observed that the number of iTALK-predicted LR interactions dropped from 108 (the lowest for mouse skin sample) in mouse skin sample to 44 (mouse cortex sample). At the same time, ICELLNET, NicheNet and NATMI still predicted over one hundred LR interactions in the mouse cortex sample.

Hence, when applied to scRNA-Seq samples, the current LR-based CCI tools can provide insight into the cell communications at the single-cell resolution. Although different tools are not often consistent, predictions from some of these tools such as iTALK, ICELLNET, and NATMI have shown good overlap with each other. They also predicted a larger percentage of well-annotated LR pairs. It would be practical for specific biological applications to run multiple tools to generate a consensus before biological validation. However, there are a few caveats to their usage and interpretation. First, the current LR annotation is incomplete. Even though there is a long list of LR pairs in various resources, many of them are computational predictions, e.g., from PPI databases, that might contain false positives. Secondly, most tools have specific details about calculating significance scores, e.g., p-values, and defining a threshold to filter confident LR pairs [27, 29]. These details need to be considered to interpret the predictions correctly. Thirdly, the cell type annotation or classification can affect the final LR-based CCI inference. Carefully selecting parameters in the cell clustering is required for a more accurate prediction. How to achieve accurate cell type classification from scRNA-Seq data is still an active topic for computational method development [94, 97, 98]. Finally, with the rapidly developing spatial transcriptomics technologies [99], methods have been developed to integrate spatial transcriptomics data and scRNA-Seq data [100, 101]. Such integrated data have demonstrated the promise to understand cell subpopulations and would hold promise for improved CCI inference in the near future [75, 79].

## Authors’ contributions

H.H. and X.L. conceived the idea. S.W. and H.Z. processed the data. S.W., X.L. and H.H. implemented the idea and generated results. S.W., J.S.C., J.K.L., X.L. and H.H. analyzed the results and contributed to the writing of the manuscript. All authors read and approved the final manuscript.

