## Supplementary Figures and Tables for "A Systematic Evaluation of the Computational Tools for Ligand-receptor-based Cell-Cell Interaction Inference"

**Supplementary Table 1.** LR pairs from 23 literatures.

| Author | Organism | Number of LR pairs | Number of one-to-one LR pairs(With unified gene symbol) | Article |
|---|---|---|---|---|
| Ding et al. (2016) | Mouse | 561 | 561 | A cell-type-resolved liver proteome |
| Yuzwa et al. (2016) | Mouse | 949 | 949 | Proneurogenic ligands defined by modeling developing cortex growth factor communication networks |
| Skelly et al. (2018) | Mouse | 2,009 | 1,863 | Single-cell transcriptional profiling reveals cellular diversity and intercommunication in the mouse heart |
| Sheikh et al. (2019) | Mouse | 1,710 | 1,621 | Systematic identification of cell-cell communication networks in the developing brain |
| Baccin et al. (2020) | Mouse | 721 | 2,251 | Combined single-cell and spatial transcriptomics reveal the molecular, cellular and spatial bone marrow niche organization |
| Cain et al. (2020) | Mouse | 2,356 | 2,356 | Quantitative single-cell interactomes in normal and virus-infected mouse lungs |
| Jin et al. (2020) | Mouse | 2,021 | 2,066 | Inference and analysis of cell-cell communication using CellChat |
| Shao et al. (2020) | Mouse | 2,033 | 1,964 | CellTalkDB: a manually curated database of ligand–receptor interactions in humans and mice |
| Kirouac et al. (2010) | Human | 232 | 565 | Dynamic interaction networks in a hierarchically organized tissue |
| Qiao et al. (2014) | Human | 933 | 923 | Intercellular network structure and regulatory motifs in the human hematopoietic system |
| Choi et al. (2015) | Human | 1,427 | 1,415 | Transcriptome Analysis of Individual Stromal Cell Populations Identifies Stroma-Tumor Crosstalk in Mouse Lung Cancer Model |
| Ramilowski et al. (2015) | Human | 2,422 | 2,422 | A draft network of ligand–receptor-mediated multicellular signalling in human |
| Pavlicev et al. (2017) | Human | 1,006 | 1,001 | Single-cell transcriptomics of the human placenta: inferring the cell communication network of the maternal-fetal interface |
| Wang et al. (2019) | Human | 2,649 | 2,587 | iTALK: an R Package to Characterize and Illustrate Intercellular Communication |
| Ximerakis et al. (2019) & BaderLab | Human | 115,900 | 23,121 | Single-cell transcriptomic profiling of the aging mouse brain |

| | | | | |
|---|---|---|---|---|
| Browaeys et al. (2019) | Human | 12,659 | 12,019 | NicheNet: Modeling Intercellular Communication by Linking Ligands to Target Genes |
| Noël et al. (2020) | Human | 380 | 456 | ICELLNET: a transcriptome-based framework to dissect intercellular communication |
| Cabello-Aguilar et al. (2020) | Human | 3,251 | 3,239 | SingleCellSignalR: inference of intercellular networks from single-cell transcriptomics |
| Jin et al. (2020) | Human | 2,005 | 2,052 | Inference and analysis of cell-cell communication using CellChat |
| Hou et al. (2020) | Human | 2,293 | 2,293 | Predicting cell-to-cell communication networks using NATMI |
| Shao et al. (2020) | Human | 3,398 | 3,398 | CellTalkDB: a manually curated database of ligand–receptor interactions in humans and mice |
| Chenf et al.(2020) | Human | 2,557 | 2,557 | Inferring microenvironmental regulation of gene expression from single-cell RNA sequencing data using scMLnet with an application to COVID-19 |
| Turei et al. (2021) | Human | 74,909 | 69,323 | Integrated intra- and intercellular signaling knowledge for multicellular omics analysis |

**Supplementary Table 2.** Robustness test results on the embryonic mouse skin dataset.

| Tool | 70% (Precision, Recall, F1) | 80% (Precision, Recall, F1) | 90% (Precision, Recall, F1) |
|---|---|---|---|
| CellChat | (0.885, 0.7887, 0.8113) | (0.9099, 0.8108, 0.8398) | (0.9347, 0.8409, 0.8695) |
| CellPhoneDB | (0.9316, 0.3963, 0.4355) | (0.9311, 0.3927, 0.4315) | (0.9435, 0.3997, 0.442) |
| iTALK | (0.9528, 0.9304, 0.9411) | (0.9605, 0.9385, 0.949) | (0.9671, 0.945, 0.9555) |
| ICELLNET | (0.9732, 0.8377, 0.8953) | (0.9733, 0.861, 0.9107) | (0.9797, 0.8891, 0.9289) |
| NicheNet | (0.8967, 0.8495, 0.8669) | (0.915, 0.8725, 0.8903) | (0.938, 0.8914, 0.9114) |
| NATMI | (0.9866, 0.8878, 0.932) | (0.9859, 0.9001, 0.9385) | (0.985, 0.9087, 0.9425) |
| PyMINEr | (0.7759, 0.6462, 0.6977) | (0.7902, 0.6898, 0.7298) | (0.802, 0.7307, 0.7579) |
| scMLnet | (0.6664, 0.5182, 0.5545) | (0.686, 0.5461, 0.5771) | (0.7142, 0.5913, 0.6182) |
| SingleCellSignalR | (0.7772, 0.667, 0.6665) | (0.7293, 0.6028, 0.5998) | (0.7732, 0.656, 0.6581) |

**Supplementary Table 3.** Robustness test results on the mouse cerebral cortex dataset.

| | 70% (Precision, Recall, F1) | 80% (Precision, Recall, F1) | 90% (Precision, Recall, F1) |
|---|---|---|---|
| CellChat | (0.8297, 0.7826, 0.7907) | (0.877, 0.7921, 0.8133) | (0.9034, 0.8086, 0.839) |
| CellPhoneDB | (0.8922, 0.3792, 0.4262) | (0.9138, 0.3763, 0.4281) | (0.9116, 0.376, 0.4286) |
| iTALK | (0.9314, 0.8685, 0.8967) | (0.9362, 0.8755, 0.9026) | (0.9387, 0.8833, 0.908) |
| ICELLNET | (0.9947, 0.9006, 0.9442) | (0.9963, 0.9257, 0.9586) | (0.9963, 0.912, 0.9511) |
| NicheNet | (0.8719, 0.8016, 0.8256) | (0.9075, 0.8274, 0.8569) | (0.9224, 0.8391, 0.8729) |
| NATMI | (0.9995, 0.9289, 0.9618) | (0.9995, 0.9401, 0.9679) | (0.9994, 0.9522, 0.9744) |
| PyMINEr | (0.7915, 0.6184, 0.683) | (0.7954, 0.6494, 0.7037) | (0.8256, 0.7051, 0.7493) |
| scMLnet | (0.6423, 0.4252, 0.4775) | (0.679, 0.4753, 0.5264) | (0.6781, 0.486, 0.5343) |
| SingleCellSignalR | (0.6043, 0.6951, 0.5679) | (0.6244, 0.6704, 0.5721) | (0.6189, 0.6765, 0.5834) |

**Supplementary Table 4.** Robustness test results on the mouse spine cord injury dataset.

| | 70% (Precision, Recall, F1) | 80% (Precision, Recall, F1) | 90% (Precision, Recall, F1) |
|---|---|---|---|
| CellChat | (0.9098, 0.8009, 0.8289) | (0.9255, 0.8108, 0.8416) | (0.9411, 0.8157, 0.8531) |
| CellPhoneDB | (0.9249, 0.4026, 0.436) | (0.9258, 0.421, 0.4485) | (0.9385, 0.4186, 0.4518) |
| iTALK | (0.942, 0.8902, 0.9144) | (0.952, 0.9098, 0.9295) | (0.9563, 0.9201, 0.937) |
| ICELLNET | (0.8403, 0.8092, 0.8197) | (0.8382, 0.8388, 0.8342) | (0.8423, 0.8379, 0.8343) |
| NicheNet | (0.8642, 0.8488, 0.8489) | (0.8937, 0.8683, 0.8752) | (0.8999, 0.8456, 0.8645) |
| NATMI | (0.9829, 0.8278, 0.8927) | (0.9817, 0.858, 0.9105) | (0.981, 0.8726, 0.9187) |
| PyMINEr | (0.7859, 0.6277, 0.6855) | (0.7988, 0.6819, 0.7245) | (0.8245, 0.7267, 0.7623) |
| scMLnet | (0.5489, 0.4234, 0.4407) | (0.5686, 0.4569, 0.4748) | (0.5922, 0.4888, 0.502) |
| SingleCellSignalR | (0.6837, 0.5353, 0.5435) | (0.7305, 0.563, 0.5812) | (0.7408, 0.5754, 0.5979) |

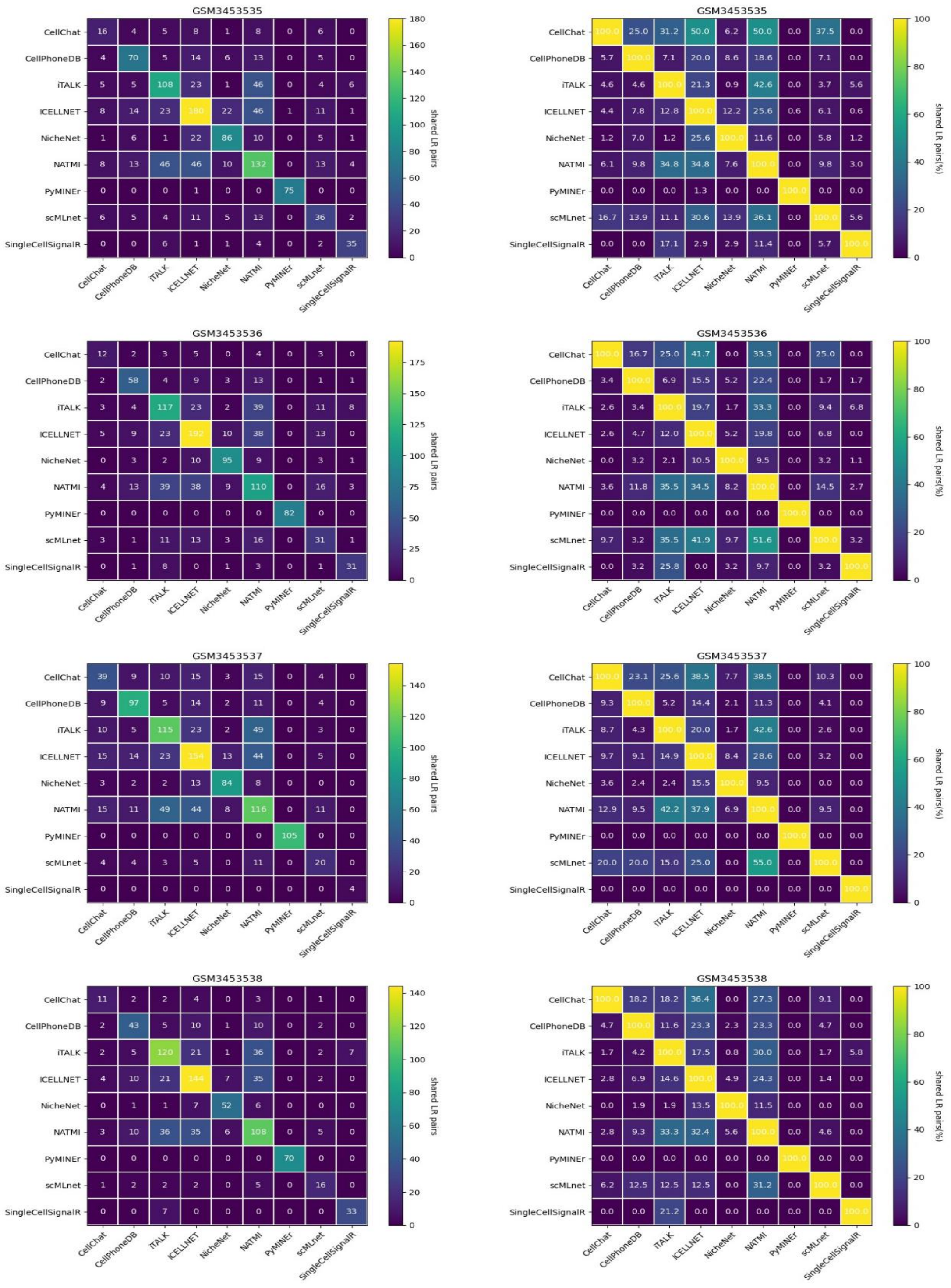

**Supplementary Figure 1.** The numbers and percentages of shared LR interaction pairs from FIB-A to FIB-B cell types in embryonic mouse skin samples.

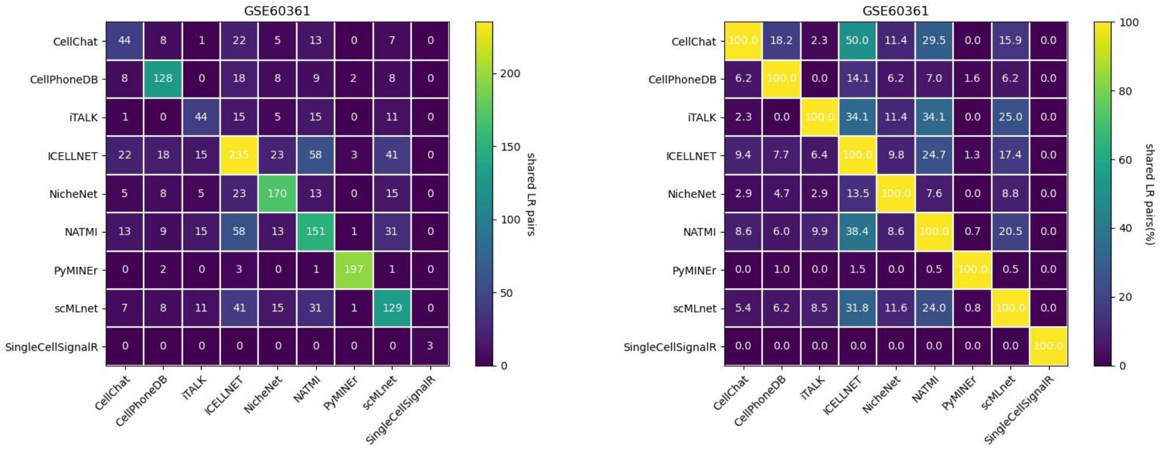


**Supplementary Figure 2.** The numbers and percentages of shared LR interaction pairs from pyramidal CA1 to oligodendrocytes cell types in mouse cerebral cortex.

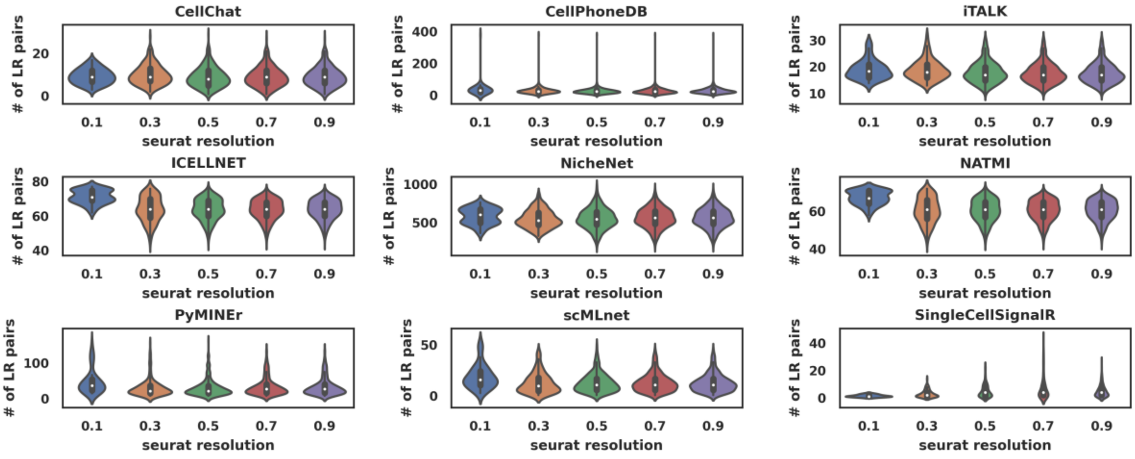


**Supplementary Figure 3.** The overall number of LR pairs across all cell types in mouse cerebral cortex under different Seurat resolution settings.

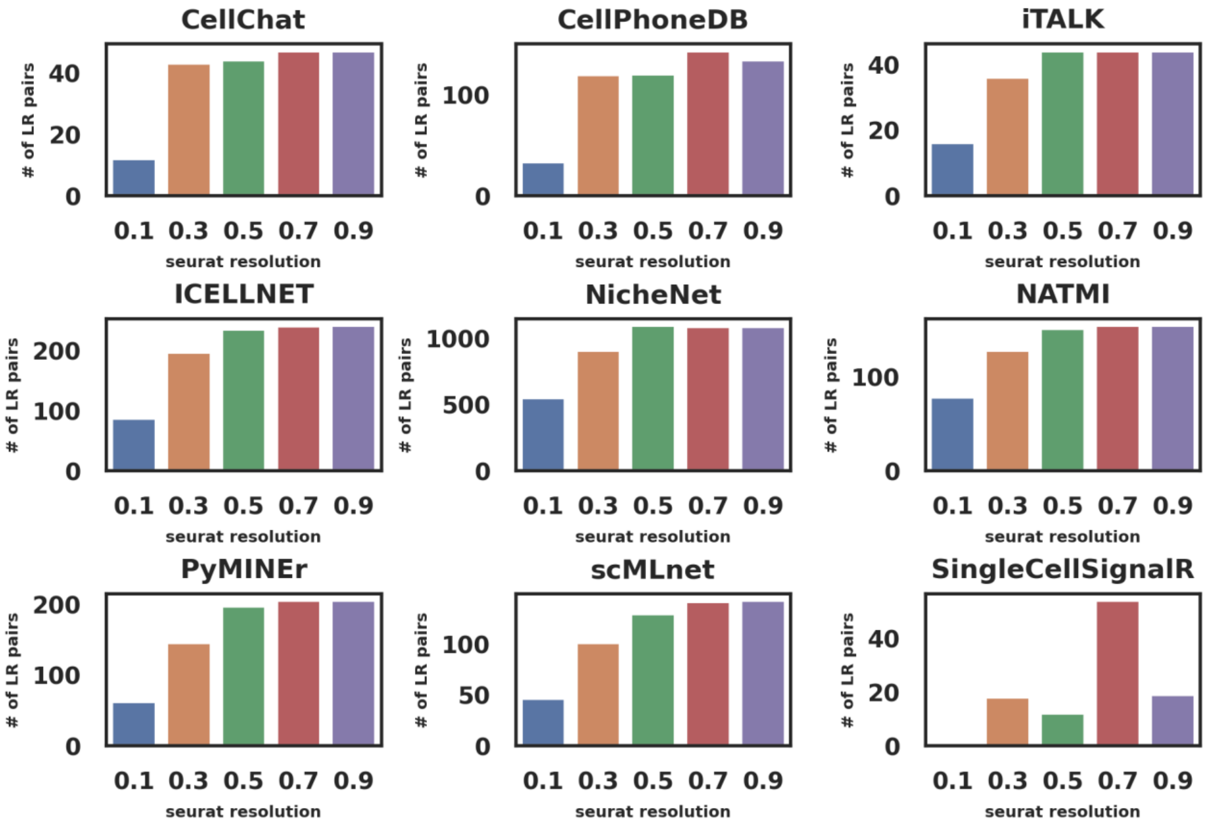


**Supplementary Figure 4.** The number of LR pairs between pyramidal CA1 and oligodendrocytes in mouse cerebral cortex under different Seurat resolution settings.

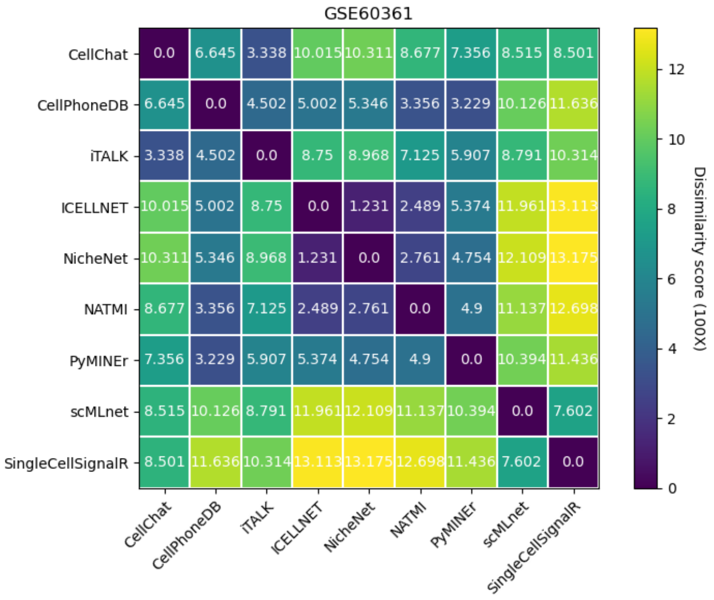


**Supplementary Figure 5.** The CCI network dissimilarity scores between different tools corresponding to the four samples in the mouse cerebral cortex dataset.

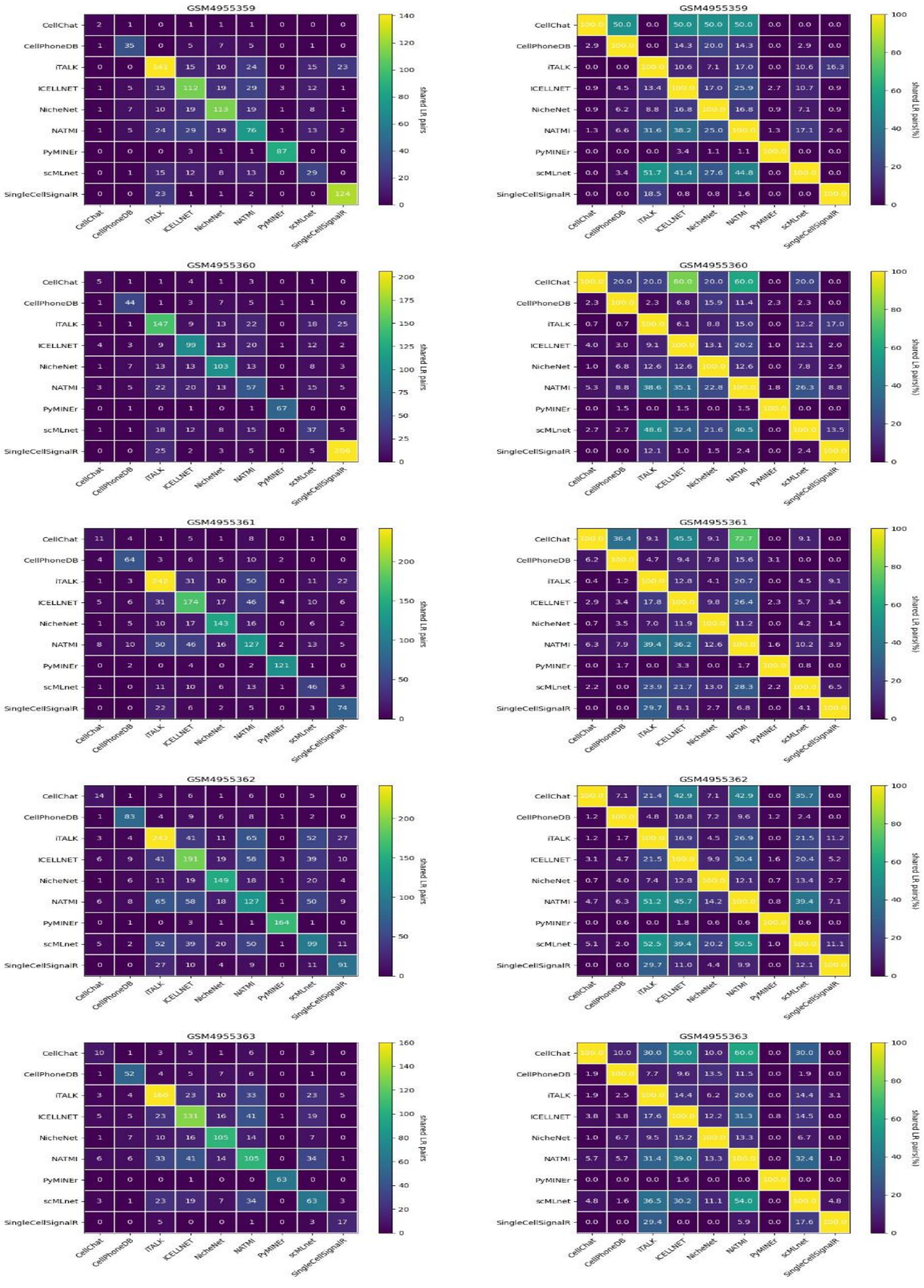
GSM4955359
GSM4955360
GSM4955361
GSM4955362
GSM4955363
CellChat
CellPhoneDB
iTALK
ICELLNET
NicheNet
NATMI
PyMINEr
scMLnet
SingleCellSignalR
shared LR pairs
shared LR pairs(%)

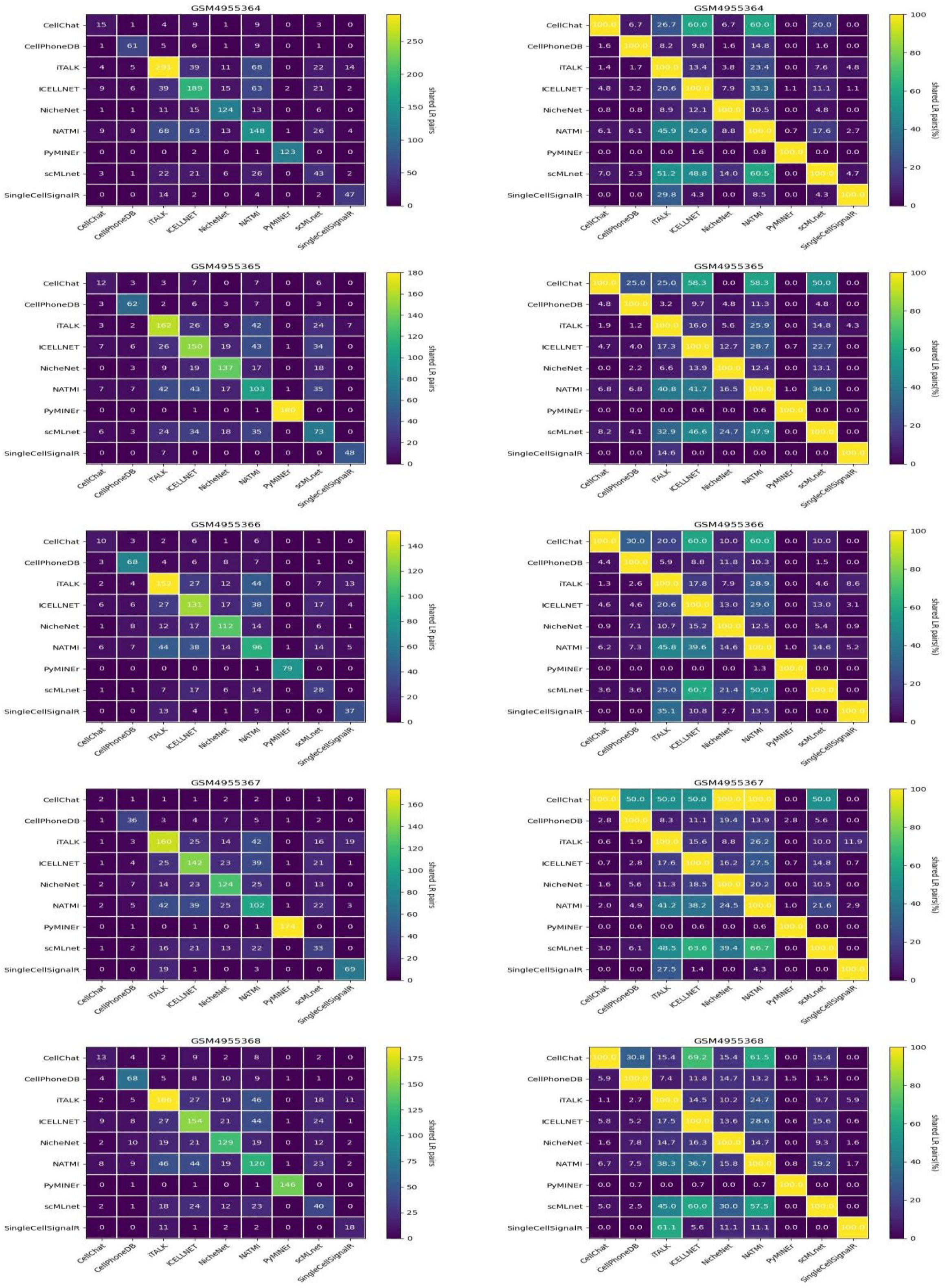


**Supplementary Figure 6.** The numbers and percentages of shared LR interaction pairs from microglia to endothelial cell types in mouse spine cord injury dataset.

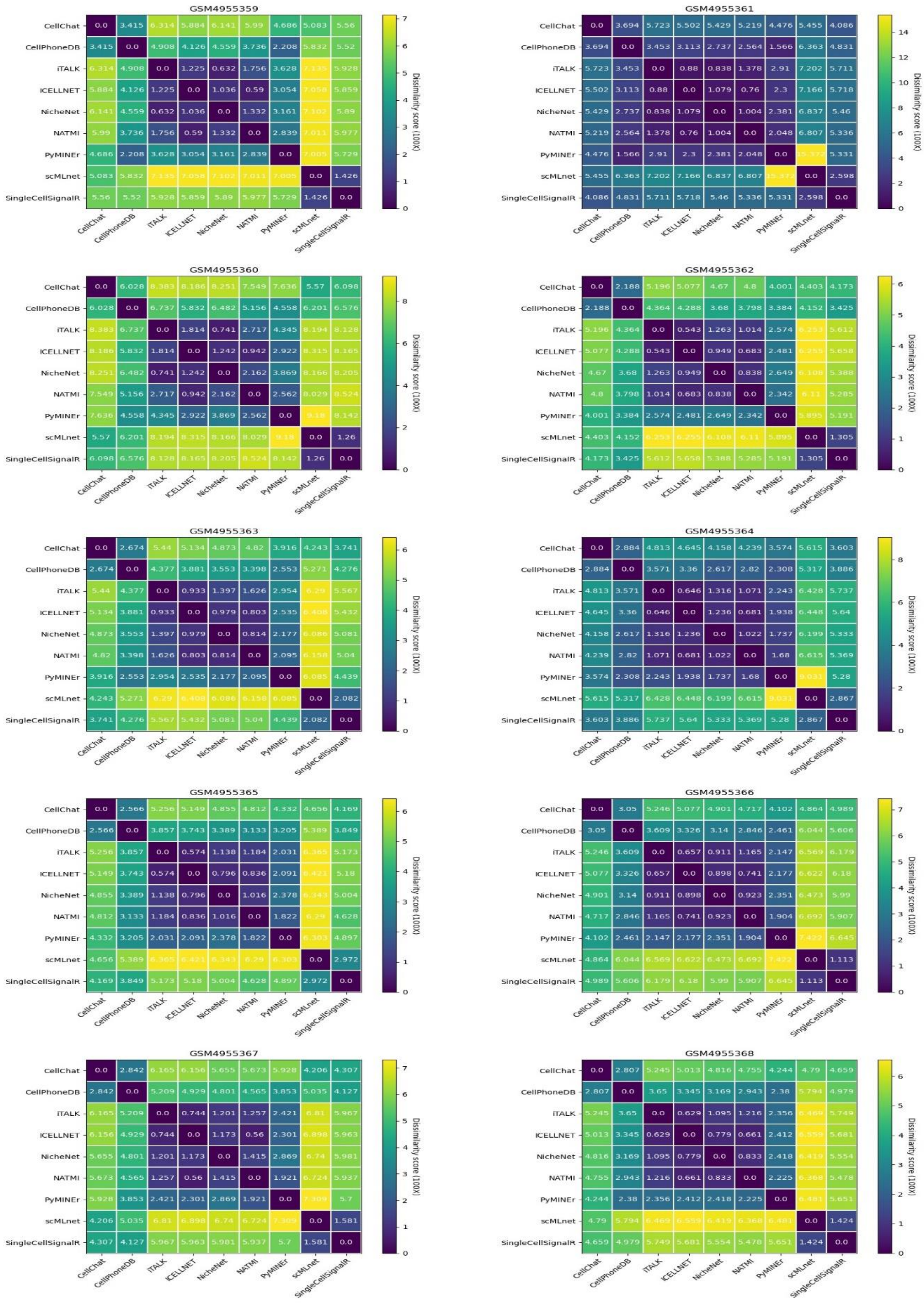


**Supplementary Figure 7.** The CCI network dissimilarity scores between different tools corresponding to the ten samples in the mouse spine cord injury dataset.
